# Neuronal cell line expressing full-length mutant huntingtin displays alteration of proteasome activity

**DOI:** 10.64898/2026.01.15.699723

**Authors:** N.N Gotmanova, T.V Bobik, A.A Ezhov, A.A Valyaeva, M.I Zvereva, M.P Rubtsova, A.V Bacheva

**Affiliations:** Department of Chemistry, Lomonosov Moscow State University, Leninskie Gory 1,3 119991 Moscow, Russia; Shemyakin-Ovchinnikov Institute of Bioorganic Chemistry, Russian Academy of Sciences, Miklukho-Maklaya str., 16/10, 117997 Moscow, Russia; Faculty of Physics, Lomonosov Moscow State University, Leninskie Gory 1,2 119991 Moscow, Russia; Belozersky Institute of Physico-Chemical Biology, Lomonosov Moscow State University, Moscow 119234, Russia; School of Bioengineering and Bioinformatics, Lomonosov Moscow State University, Moscow 119234, Russia

## Abstract

Polyglutamine diseases are incurable genetic neurodegenerative disorders characterized by the accumulation of extended polyglutamine fragments-containing mutant proteins, which are prone to the formation of poorly soluble aggregates. The adequate cellular model is crucial in uncovering the pathological mechanisms responsible for neurotoxicity in HD, screening for therapeutic molecules and elucidation of the molecular mechanisms impacted by particular compound. In the present study, genetic constructs based on the Sleeping Beauty system were created for the stable inducible expression of full-length normal and mutant huntingtin (Htt) in eukaryotic cells. These constructs were then employed to develop model neuronal cells using Neuro-2a cell line. The expression of Htt as well as the accumulation of Htt immunopositive intracellular aggregates (most characteristic features of HD) was demonstrated, and these aggregates showed colocalization with the proteasome. The activation of the proteasome, as well as changes in the expression of proteasome regulators, components of the autophagy system, and the neuronal proteases cathepsins B and D, reflect the versatility of cellular responses to the mutant pathological forms of Htt.

## Introduction

Huntington’s disease (HD), a prominent example of polyglutamine proteinopathies, is associated with the selective death of GABA-ergic neurons in the striatum of the brain and is caused by a pathological mutation in the *HTT* gene encoding huntingtin (Htt), a ubiquitously expressed approximately 350 kDa protein. Wild-type *HTT* alleles contain up to 35 continuous CAG repeats in the first exon, thereby expressing into a normal polyglutamine (polyQ) tract, while mutant alleles are characterized by 36 or more repeats. Huntingtin coordinates several vital processes in the cytoplasm and nuclei of neurons [1, 2]. It is assumed that the mutant huntingtin (mHtt) is unable to perform its physiological role and can also acquire atypical neurotoxic functions [3–6]. Abnormally long polyQ stretch is the major factor contributing to mHtt pathological aggregates formation that lead to neuronal dysfunction and death. These aggregates often contain ubiquitin and proteasome subunits [7–11]. General mHtt degradation mechanisms include ubiquitin-proteasome system, chaperones and autophagy [12–15], but mHtt proteolysis by other cellular proteases has also been described [16–19]. However, the molecular events connecting how mHtt aggregation triggers proteostasis imbalance and the origins of mHtt aggregated forms proteotoxicity have not been studied in detail. In this context, a comprehensive study of huntingtin variants proteostasis in various model systems remains relevant.

Recent studies of mHtt proteostasis in eukaryotic cell lines have focused on huntingtin exon 1 (normal length is 82 aa) [20] with 70-150 glutamine residues in polyQ tract [21–23], since mHtt N-terminal proteolysis products are the most toxic for proteostatic balance [3]. I*n vitro* HD cell models are irreplaceable for reproducing mHtt pathogenic effects from whole organisms. The rat neuroblastoma-glioma model NG108-15 with inducible expression of full-length mHttQ116 demonstrates time-dependent formation of cytoplasmic and nuclear Htt-positive inclusions, the latter being colocalized with the proteasome regulator PA700 [24]. In the HEK293 Tet-Off system expressing huntingtin exon 1 (Q20, Q51 or Q83), the number, size and toxicity of ubiquitinated Htt-immunopositive inclusion bodies are polyQ length-dependent. Htt-Q83 aggregates also showed colocalization with 20S, 11S, and 19S proteasome subunits, and proteasome inhibition dramatically enhance mHtt fragments aggregation and cytotoxicity [25]. However, in this model mHtt aggregates had predominantly perinuclear localization. Among cellular models HD-associated proteostasis of polyQ fragments has been investigated predominantly in non-neuronal cells or neuronal precursors, including yeast models [26], HEK293 [27] and HeLa cells [28], insect cells [29, 30], and others [31, 32]. Plant BY-2 tobacco cells stably expressing EGFP-tagged Htt exon 1 (Q23/Q52) could serve as a platform for screening compounds with anti-aggregation activity, although this model reproduces only the mHtt fragments aggregation [32]. The HEK293T model with transient transfection of mHtt-128Q N-terminal fragments (771 or 1955 aa) [33] demonstrated that the mHtt aggregates toxicity is independent of their intracellular localization, but associated with the frequency of aggregate formation. Conversely, prior studies on HD transgenic mouse models have demonstrated that intranuclear mHtt deposits exhibited the most destroying neurotoxic effects [34]. The formation rate, quantity and toxicity of mHtt aggregates remains time- and polyQ length-dependent in each three different cell lines (COS-7, PC12 and SH-SY5Y), transiently transfected with huntingtin exon 1 (23 or 43-74Q) [35]. The aggregation patterns may also depend on the whole mHtt N-terminal fragments length (polyQ-independent aggregation mechanisms for fragments longer than 170 aa) and its expression level [36]. In general, transient expression of truncated/full-length Htt could result in side effects on transcriptomic profile and cell metabolism, while expression of huntingtin N-terminal fragments is unable to recapitulate the full range of typical HD disturbances. HD models from reprogrammed patient fibroblasts or primary neuronal cultures require high time costs and resources, and resultant cell populations usually comprise diverse neuron types, depending on the neurotransmitter they generate. Consequently, we have chosen to develop a new cellular model for Huntington’s disease based on neuronal cell line Neuro-2a [37, 38] with stable expression of full-length normal or mutant human huntingtin. The primary objective of the present study was to investigate the full-size mHtt proteotoxicity, which can be mediated not only by N-terminal fragments [39]. Moreover, full-length mHtt protein is required for accurate reproduction of the pathogenic effects, since normal Htt serves as a molecular scaffold for assembling protein complexes and regulating distinct intracellular pathways [1]. The present study aimed to explore the roles of cellular proteolysis systems, specifically the proteasome system, lysosomes, and autophagy, in the HD cellular model.

## Materials and methods

### Plasmids construction

The original pSB*tet*-Neo vector was from Addgene (#60509). After pSB*tet*-Neo PCR linearization (using Q5® High-Fidelity DNA Polymerase, New England Biolabs, USA) with introduction of MluI and NotI restriction sites (forward primer with NotI site: 5′-AATAGCGGCCGCGCTTCCATCGATAGACATGATAAGATAC-3′, reverse primer with MluI site: 5′-TATTACGCGTCAGAGGCCTTTCGAGGGTAGG-3′; restriction sites are underlined) as well as restriction of the pCI-HttQ15 and pCI-HttQ138 plasmids [40] at MluI and NotI sites, the resulted fragments were ligated (insertion:vector = 1:3 in molar ratio) and transformation of competent *E. coli* XL1-Blue cells with the obtained ligase mixture was carried out. Bacterial colonies containing pSBtet-Neo-HttQ15/Q138 construct were identified by PCR using primers flanking C-terminal region of Htt gene (forward primer: 5′CCACCTGCTGACTACAAAGACCAT-3′) and the start of the pSB*tet*-Neo vector promoter region (reverse primer: 5′-CCATTGAGTAATTCCAGAGCGCCG-3′). As expected, the length of observed PCR products was about 1100 b.p. (Supplementary, Fig. S1). Isolation of plasmid DNA from *E. coli* cells was performed using Plasmid Miniprep kit (Evrogen, Russia) according to the manufacturer’s instructions. The accuracy of the insertion nucleotide sequence in the selected clones was confirmed on MinION sequenator using the Oxford Nanopore R9.4.1 flow cell (Oxford Nanopore technology, UK). ONT reads quality control was performed using NanoPlot (version 1.41.6) [41]. Reads were grouped into two categories (Q15 and Q138) according to the pSBtet-Neo-HttQ15/Q138 construct utilized. Initially, reads were aligned to the linearized reference plasmid sequence using minimap2 (version 2.26-r1175) [42]. For subsequent analysis only reads that covered the N-terminal region of the Htt gene (nucleotides from 1 to 147) were retained. The number of CAG repeats in polyQ tract was estimated using pipeline consisting of LAST aligner (version 1454) [43] and tandem-genotypes (version 1.9.0) [44] (https://github.com/mcfrith/tandem-genotypes). The visualization of reads alignment on reference plasmid maps was carried out using Galaxy (https://usegalaxy.org/) and IGV 2.13.1 software. Code for replicating ONT reads analysis and figures was deposited to GitHub (https://github.com/kirushka/nanopore_huntingtin).

### Neuro-2a culture and transfection

Neuro-2a (mouse neuroblastoma) cells wase kindly provided by Dr. A.A. Kudriaeva (Shemyakin-Ovchinnikov Institute of Bioorganic Chemistry, Russian Academy of Sciences, Moscow, Russia). The cells were originally obtained from shared research facility “Vertebrate cell culture collection” of the Institute of Cytology, Russian Academy of Sciences. Cells were cultured in sterile CO_2_-incubator (37°С, humidified atmosphere, 5% CO_2_) on DMEM-F12 media containing 10% FCS, 1% Pen-Strep, 2 mM *L*-glutamine and 4,5 g/L glucose (Gibco, USA). When cells were reached 80% confluence, co-transfection was performed in Opti-MEM media (Gibco, USA) using Lipofectamine 3000 transfection kit (Invitrogen, USA) according to the manufacturer’s instructions. 1 μg of total plasmid DNA (950 ng pSBtet-Neo-HttQ15/HttQ138 + 50 ng SB100x transposase coding vector pCMV(CAT)T7-SB100 (Addgene, Plasmid #34879) was applied to cells in one co-transfection experiment.

### Generation of monoclonal transgenic Neuro-2a cell lines expressing normal or mutant huntingtin variants

After co-transfection, the cell culture medium was changed to DMEM-F12 containing 1 mg/ml antibiotic G418 (Sigma-Aldrich, USA). Parallel control experiments included: cells with co-transfection and without G418 (positive control) and cells without co-transfection and with G418 (negative control). Then G418 was added for 10-12 days until 100% cell death was achieved in the negative control. A portion of the resulting polyclonal culture was then transferred to a 96-well culture plate, seeding 1, 5 and 10 cells/ml in each 32 wells. Selection was carried out for 10-14 days. As a result, 4 candidate Neuro-2a monoclones with each huntingtin gene variant were selected.

### qPCR determination of HttQ15/HttQ138 transgenes expression levels in Neuro-2a transgenic cells

Neuro-2a poly- and monoclones with HttQ15/HttQ138 stable transfection, as well as cells transfected by pSB*tet*-Neo «empty» construct without gene insertion, were induced by 1 μg/ml doxycycline; in the control transgenes set there was no doxycycline induction. After 2 days of induction, Neuro-2a cultures were lysed with 500 μl/5,0·10^5^ cells QIAzol reagent (QIAGEN, Germany). After phenol-chloroform protein extraction and aqueous phase separation, RNA was precipitated by absolute isopropanol, washed by 75% ethanol and air-dried. Obtained RNA precipitates were dissolved in milliQ water and put into ice. Total RNA concentrations were determined by 260 nm absorption using NanoDrop-2000 spectrophotometer (Thermo Scientific, USA). The total RNA yield was 50-80 μg/5,0·10^5^ cells. Then, 1 μg of each RNA sample was treated with 10 units of DNAse I (Thermo Fisher Scientific, USA) to remove any DNA traces. Total cDNA first chain synthesis was carried out using Magnus reverse transcriptase (Evrogene, Russia) and a random decanucleotide primer according to the manufacturer’s instructions. Parallel control reactions included all the required components except RNA matrix (negative control).

qPCR transgenes expression analysis was performed using 5X qPCRmix-HS SYBR (Evrogene, Russia), forward (5′-AATGGTGCCCCTCGGAGTTTGC-3′) and reverse (5′-CTGCATTTCTGAGGCCGAACCAGG-3′) primers for Htt gene fragment amplification, as well as forward (5′-TGCACCACCAACTGCTTAGC-3′) and reverse (5′-GGCATGGACTGTGGTCATGAG-3′) primers for GAPDH gene fragment amplification (normalization household gene). Each reaction mixture contained 150 ng of appropriate cDNA matrix. Parallel control experiments included the cDNA matrix replacement with negative reverse transcription control or milliQ water (negative PCR control). Each reaction (except negative controls) was performed in three technical replicates. Amplification was performed on a CFX96 real-time system (Bio-Rad, USA) using CFX Manager™ software (Bio-Rad, USA).

The relative transgene expression increase compared to the absence of induction was expressed by 2^−ΔΔCt^ value [45], where C_t_ — threshold cycle number, ΔC_t_ = C_t_(HttQ15/Q138, without induction) – C_t_(GAPDH), ΔΔC_t_ = ΔC_t_(sample with induction) – ΔC_t_(sample without induction). The PCR products were additionally analyzed by agarose gel electrophoresis.

### Determination of HttQ15/HttQ138 transgenes copy number

Poly-/monoclonal Neuro-2a cells, as well as control cells (transfected with «empty» pSB*tet*-Neo vector and non-transfected) were lysed with QIAzol reagent, as described above. After phenol-chloroform extraction, the aqueous phase was removed from the samples and DNA was extracted with 100% ethanol. Next, the precipitate was washed twice with 0,1 M sodium citrate in 10% ethanol, resuspended in 75% ethanol, centrifugated, dried and dissolved in 8 mM NaOH, and pH was adjusted to 7,5 with a HEPES buffer. Genomic DNA concentrations were determined by 260 nm absorption using NanoDrop spectrophotometer. DNA yield was 50-100 μg/5.0·10^5^ cells.

The transgene copy number determination was carried out by the qPCR, as described in the previous section, using the same primers. Each reaction mixture contained 10 ng of appropriate cDNA matrix. The estimation of transgene inserts number was carried out according to ΔΔC_t_ method, as described above. The GAPDH-normalized Htt gene copies number in Neuro-2a cells with «empty» pSB*tet*-Neo construct was taken as 1.

### Immunofluorescence microscopy

Monoclonal (Q15m/Q138m) and polyclonal (Q15p/Q138p) Neuro-2a lines, as well as control cells ((−) and −SB) were seeded on poly-*L*-lysine covered 10 mm slide glasses 24–48 h prior to imaging. Cells were washed with PBS, fixed with 3,7% paraformaldehyde for 5 min, then permeabilized with 1% Triton X-100 for 15 min, blocked with 1% BSA in PBST and then incubated with primary antibodies: anti-Htt (1:50, Elabscience, USA), anti-α1, 2, 3, 5, 6, 7 (1:100, Enzo Life Sciences, USA); and subsequently incubated with secondary antibodies: Alexa Fluor 488-conjugated goat-anti-mouse (1:150, Invitrogen, USA), Alexa Fluor 568-conjugated donkey-anti-rabbit (1:150, Invitrogen, USA). Cell nuclei were stained with 1 μg/ml DAPI. The fluorescent staining was analyzed on motorized inverted microscope Olympus IX81 (Japan). Fluorescent images were obtained by the filter cubes U-MWU2, U-MNB2 and U-MWG2 (all — Olympus, Japan) in combination with Xe short arc lamp as the excitation light source. Dry objective lens UPLSAPO40X2 (NA = 0.95) and oil immersion objective lens UPLSAPO60XO (NA = 1.35) (both — Olympus, Japan) were used. The images were recorded using cooled monochrome CCD camera XM10 and color CCD camera DP26 (both — Olympus, Japan) and processed using software ImageJ version 1.54g (National Institutes of Health, USA). Colocalization quantitative characteristics were estimated with the same ImageJ software.

### Filter trap assay

Cells were detached with 0,5% (1x) trypsin-EDTA solution (Gibco, USA), collected, harvested by centrifugation, and washed 3 times with PBS. Then 200 μl of lysis buffer (10 mM Tris-HCl, pH 7,5, 150 mM NaCl, 2% SDS) was added to each cell sample and lysates were sonicated on Ultrasonic Processor (Cole-Parmer, USA) by 1 impulse for 5 seconds, 50 W. The total protein concentration in the cell lysates was determined using Pierce™ BCA Protein Assay Kit (Pierce, USA). The samples were diluted with a lysis buffer so that the total protein concentration in the samples was 0,4 μg/μl in a volume of 400 μl (1:1 dilution), subsequential dilutions (1:5, 1:25) by lysis buffer were made and DTT was added to each sample to the 50 mM final concentration. For samples transferring on 0,45 μm nitrocellulose membrane HyBond C Extra (Amersham, USA) Dot Blot 96 system (Biometra, Germany) with vacuum manifold was used. 100 μl of each sample was subjected to appropriate Dot Blot 96 system well on membrane pre-equilibrated with wash buffer (10 mM Tris-HCl, pH 7,5, 150 mM NaCl, 0,1% SDS) under mild vacuum. Each well containing sample was washed three times by 100 μl of wash buffer. The membrane was washed three times by PBS, blocked with 5% BSA in PBST buffer, then stained by 0,5% (w/v) Ponceau S dissolved in 1% (v/v) acetic acid and photographed on ChemiDoc XRS+ (Bio-Rad, USA) imaging system. After Ponceau S washing, membrane was incubated with primary antibodies: anti-Htt or HRP-conjugated anti-FLAG epitope (1:1000, Merck Millipore, USA); and subsequently incubated with secondary antibodies: HRP-conjugated goat-anti-mouse (1:2000, Sigma-Aldrich, USA). The chemiluminescence (ECL) signal was detected using ChemiDoc XRS+ (Bio-Rad, USA) imaging system and Image Lab™ 6.0.1 software (Bio-Rad, USA).

### Proteasome and cathepsins B and D activity measurements

After 7 or 14 days of doxycycline induction (in the absence of the inducer for negative control) monoclonal (Q15m/Q138m) and polyclonal (Q15p/Q138p) Neuro-2a lines, as well as control cells ((−) and −SB) were detached with cell scraper, harvested by centrifugation, and washed 3 times with PBS. Cellular proteasome activity was measured using synthetic peptide fluorogenic substrates (Suc-LLVY-AMC, Z-LLE-AMC and Ac-RLR-AMC for chymotrypsin-like, caspase-like and trypsin-like activity detection, respectively). Briefly, 10 μl of 1,0·10^4^ cell sample suspended in PBS was added to each substrate solution (50 μM Suc-LLVY-AMC, 50 μM Z-LLE-AMC or 20 μM Ac-RLR-AMC) in 50 mM Tris-HCl, 1 mM EDTA, 0,5% Triton-X100, pH 7,5 buffer in the opaque black 96-well microplate wells (total volume of each reaction was 100 μl). To determine non-proteasome proteolytic activity (only in Q138p and SB transgenic lines), proteasome inhibitor MG132 was preliminarily added to the substrate solutions (150 nM, 3 μM or 10 μM for 100% inhibition of chymotrypsin-like, caspase-like or trypsin-like proteasome activity, respectively) and the to the control wells. The fluorescence intensity was measured on a Victor X5 2030 microplate reader (Perkin Elmer, USA) for 45 min with 20 s interval (excitation wavelength = 355 nm, emission wavelength = 460 nm, 37°С). Each experiment was carried out in three technical replicates.

The proteasome activity (per reaction volume) was calculated by the formula:

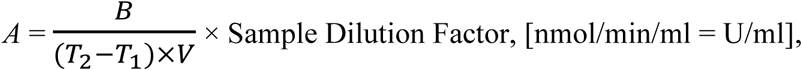

where *B* is the AMC amount from AMC standard curve (pmol), *T*_1_ is the time of the first fluorescence reading (min), *T*_2_ is the time of the second fluorescence reading (min), *V* is the sample volume added into the reaction well (μl). One unit of proteasome activity is defined as the amount of proteasome which generates 1,0 nmol of AMC per minute at 37°C. Proteasome activity for each transgenic/control Neuro-2a line in the absence of doxycycline induction was referred to 100%. Background protease activity was subtracted form activity values in each experiment. To determine the fluorescence intensity of free AMC, a calibration measurement was performed in the 5-200 μМ range of AMC concentrations. For results processing, Perkin Elmer 2030 Manager (PerkinElmer Life Sciences) and SigmaPlot 12.5 (Systat Software) software was used.

Intracellular cathepsin B activity measurement was conducted using the same approach. Z-Phe-Arg-AMC (50 μM) was used as a synthetic peptide fluorogenic substrate; the reaction buffer was 50 mM imidazole-HCl, 4 mM EDTA, 8 mM DTT, 0,5% Triton-X100, pH 4,8; the cysteine protease inhibitor E64 (100 nM) was used to 100% inhibit the activity of cathepsin B.

Cathepsin D activity was measured on cellular lysates after resuspension in lysis buffer (50 mM Tris-HCl, pH 7,5, 150 mM NaCl, 1 mM EDTA, 1 mM DTT, 0,5% Nonidet-P40) and clarification of supernatant by centrifugation. 40 μM Abz-Ala-Ala-Phe-Phe-Ala-Ala-DeD (where Abz is anthranilic acid residue, DeD is 2,4-dinitrophenylethylenediamine residue) was used as a synthetic peptide fluorogenic substrate with intramolecular fluorescence quenching; the reaction buffer was 0,1 M Na-citrate, 4 mM EDTA, 0,5% Triton-X100, pH 4,1; the protease inhibitor pepstatin A (100 nM) was used to 100% inhibit the activity of cathepsin D. The total protein concentration in the cell lysates was determined using Pierce™ BCA Protein Assay Kit (Pierce, USA). The fluorescence intensity was measured on a CLARIOstar Plus microplate reader (BMG Labtech, Germany) for 45 min with 10 s interval (excitation wavelength = 340 nm, emission wavelength = 415 nm, 37°С). For results processing, CLARIOstar Plus, MARS Data Anaysis (BMG Labtech, Germany) and SigmaPlot 12.5 (Systat Software) software was used.

### MTT viability assay

Neuro-2a cell lines were seeded into 96-well transparent plate (1,0·10^4^ cells/well), induced by 1 μg/ml doxycycline or cultured without induction. After induction period, 10 µl of MTT solution in sterile PBS (5 mg/ml) was added to each well and the cells were incubated in a sterile CO_2_ incubator (37°C, humidified atmosphere, 5% CO_2_) for 3,5 hours. Then the medium was removed from the cells, 100 µl of DMSO was added to dissolve the formazan precipitate and the plates were stirred at room temperature in the dark for 30 minutes. The absorption of the obtained samples was determined on a Victor X5 2030 microplate reader (Perkin Elmer, USA) at 590 nm wavelength. The viability of each cell line was calculated according to the formula:

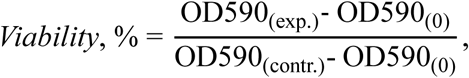

where OD590_(exp.)_ is the average optical density of the experimental sample, OD590_(contr.)_ is the average optical density of the negative control (no doxycycline induction), and OD590_(0)_ is the average background optical density (DMSO).

### Western blotting experiments

Cells were detached with 0,5% (1x) trypsin-EDTA solution (Gibco, USA), collected, harvested by centrifugation, and washed 3 times with PBS. Then cells were resuspended in lysis buffer (50 mM Tris-HCl, pH 7,5, 150 mM NaCl, 1 mM EDTA, 1 mM DTT, 0,5% Nonidet-P40, supplemented by protease inhibitor cocktail cOmplete Mini EDTA-free), frozen on −70°C/thawed 3 times and sonicated on Ultrasonic Processor (Cole-Parmer, USA) by 6 impulses for 10 seconds, 130 W. The total protein concentration in the cell lysates was determined using Pierce™ BCA Protein Assay Kit (Pierce, USA). Cell lysates were boiled for 10 min at 99°С with 1x Laemmli sample loading buffer (50 mM Tris-HCl, 10% (v/v) glycerol, 2% SDS, 1% β-mercaptoethanol, 12,5 mM EDTA, 0,02% bromophenol blue, pH 6,8) and after SDS-PAGE fractionation, were transferred on 0,2 μm nitrocellulose membrane HyBond C Extra (Amersham, USA). The membranes were blocked with 5% BSA in PBST buffer, then incubated with primary antibodies: anti-Htt (1:1000, Merck Millipore, USA), anti-α1, 2, 3, 5, 6, 7 (1:1000, Enzo Life Sciences, USA), anti-β1, anti-β2, anti-β5, anti-β1i, anti-β2i, anti-β5i (1:1000, Enzo Life Sciences, USA), anti-11Sα (1:1000, Santa Cruz, USA), anti-11Sγ (1:1000, Abcam, UK) anti-19S (1:1000, Enzo Life Sciences, USA), anti-cathepsin D (1:500, Elabscience, USA), anti-LC3B (1:500, Elabscience, USA), anti-β-actin (1:1000, Santa Cruz, USA); and subsequently incubated with secondary antibodies: HRP-conjugated goat-anti-mouse (1:2000, Sigma-Aldrich, USA), HRP-conjugated goat-anti-rabbit (1:2000, Sigma-Aldrich, USA). The chemiluminescence (ECL) signal was detected using ChemiDoc XRS+ (Bio-Rad) imaging system and Image Lab**^TM^** 6.0.1 software (Bio-Rad, USA).

### Statistical analysis

All values were obtained from three independent repeated experiments and expressed as mean ± S.D. Statistical analysis was performed using Student’s *t* test. *p* < 0,05 was considered as statistically significant.

## Results

### Generation of stable, inducible eukaryotic cell lines for HttQ15/HttQ138 expression

To create a genetic construct for the stable expression of full-length huntingtin (Htt) with normal polyglutamine tract (HttQ15) or elongated polyglutamine tract (HttQ138) in eukaryotic cells, HTTQ15 or HTTQ138 gene from pCI_HTT Q15/138 plasmid [40] was cloned into the pSB*tet*-Neo expression vector. In this vector, the gene of interest is flanked by transposon sequences (ITR, inverse terminal repeats). Following the co-transfection of eukaryotic cells with the pSB*tet*-Neo-HTTQ15/Q138 construct and the SB-100x transposase encoding vector, transgenic cells were obtained with stable doxycycline-inducible expression of HttQ15 or HttQ138 [46]. Supplementary Figures S1 and S2 illustrate the process of huntingtin genes being cloned into the pSBtet-Neo vector and the PCR screening results of *E. coli* colonies containing the required plasmids.

A number of candidate clones were selected for each of the pSB*tet*-Neo-HTTQ15/Q138 constructs based on PCR screening results. The plasmid DNA extracted from selected clones was subjected to Sanger sequencing, and then to Nanopore sequencing to confirm the total structure of the plasmid. This method is particularly well-suited for the sequencing of large constructs (∼15 000 b.p.), however, it should be noted that it is not designed to accurately estimate the number of repeats in cases where the repeats are exceedingly abundant. The analysis of the Nanopore sequencing data demonstrated that the distribution of read length exhibited a prominent peak around 15kb, which corresponds to the full-length plasmid sequencing reads. The analysis of CAG copy number was performed using reads that contained the Htt N-terminal region. The analysis revealed that the median CAG repeat length in pSB*tet*-Neo-HttQ15 samples was 14, and in the pSB*tet*-Neo-HttQ138 vector it was 100-130 repeats with a modal value of 136 CAG repeats (Fig. S3). A restriction analysis was conducted to verify the compatibility of the pSB*tet*-Neo-HttQ15/Q138 vectors with the reference sequences (Fig. S4). The observed fragments’ lengths correspond to the lengths predicted based on the location of the hydrolysis sites of BamHI and HindIII in both expression vectors.

Then Neuro-2a cell lines with stable expression of full-length HttQ15/HttQ138 forms were developed. The co-transfection of each pSB*tet*-Neo-HTTQ15/Q138 plasmids and the SB-100x transposase gene encoding vector was performed by lipofection. The G418 antibiotic (1 mg/ml) was used for selection of stable transfects as it resulted in the complete eradication of 2.0-5.0·10^5^ non-transfected Neuro-2a cells within 3 days (data not shown). The selection of stable transfects was performed by adding of G418 over a period of 10 days. The control experiments included cells without transfection, transfected with an “empty” pSB*tet*-Neo plasmid without HTT genes, and with transfection of both huntingtin variants, excluding the addition of G418. Following the complete elimination of living cells in the non-transfected control culture was attained, the selection process was extended for a further 2–3 days. The resulting polyclonal transgenic Neuro-2a cultures containing the HTTQ15 or HTTQ138 gene were then subjected to a clonal selection procedure, which produced four candidate clones for each variant of the Htt gene (Fig. 1a).

**Figure 1.**
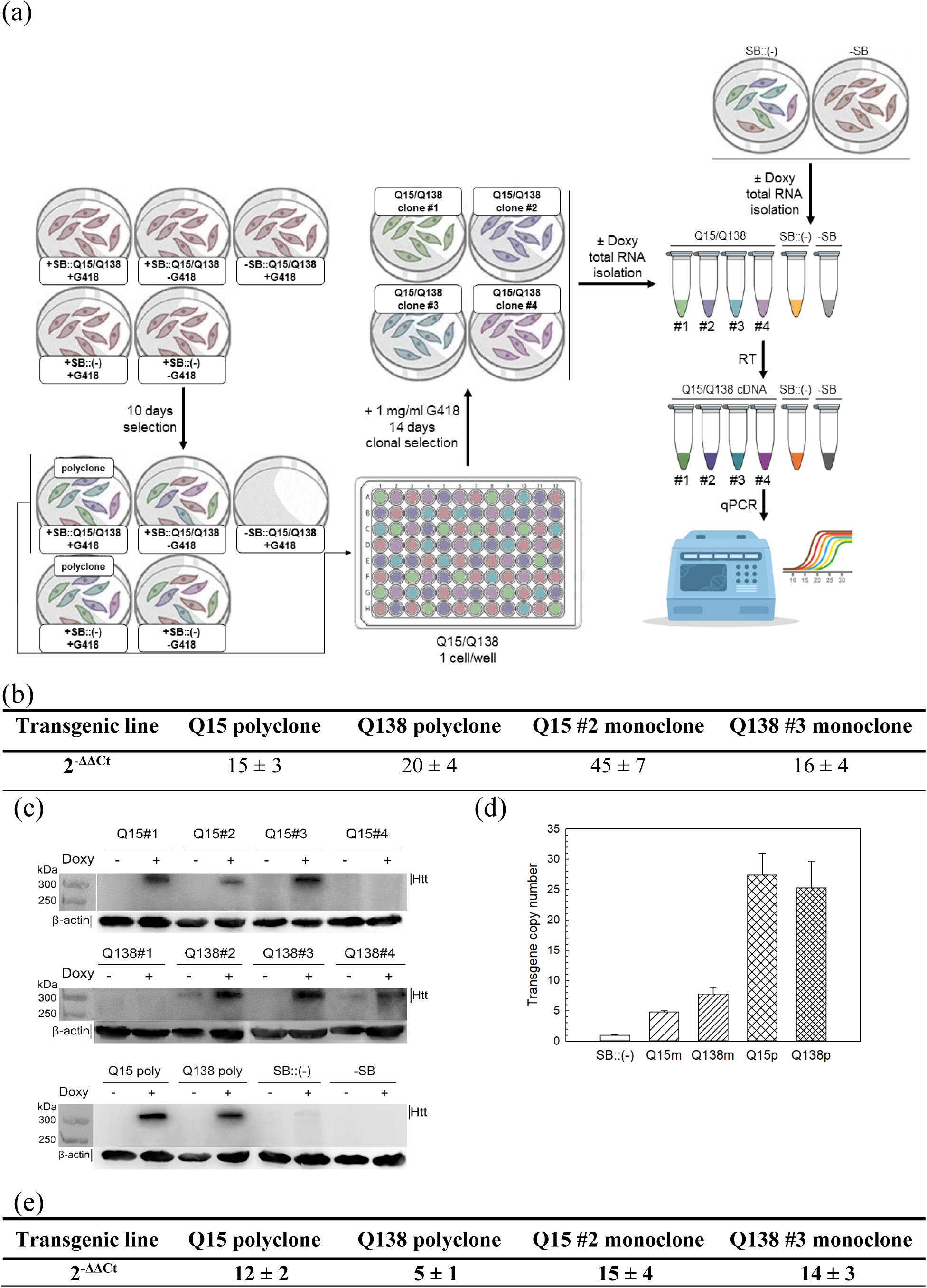
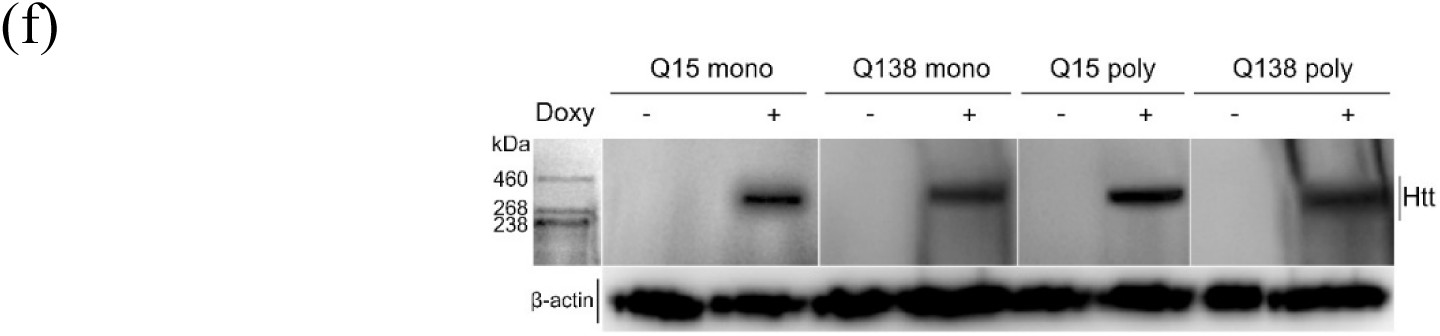
Application of the Sleeping Beauty system for the transposition of normal/mutant huntingtin genes into the genome of Neuro-2a cells allows to create HD transgenic cellular model and determine the levels of HttQ15/Q138 genes expression in obtained cell lines. a) The validation of transgenes expression by quantitative PCR (qPCR). SB::Q15/Q138 – pSB*tet*-Neo vector containing HttQ15 or HttQ138 gene, (−) – pSB*tet*-Neo vector without huntingtin gene insertion, −SB – Neuro-2a cells without transfection, RT – reverse transcription; b) 2^−ΔΔCt^ values for transgenic Neuro-2a lines with HttQ15 or HttQ138 expression; c) Western blotting of poly/monoclonal transgenic Neuro-2a cells lysates without/with doxycycline induction of HttQ15 or HttQ138 expression. Data for control Neuro-2a lines ((−), with transfection of «empty» vector, and −SB, without transfection, are represented also. Top panel – lysates of four different monoclonal transgenic Neuro-2a lines with HttQ15 expression, middle panel – same as top, but for lysates of HttQ138 monoclonal lines, bottom panel – lysates of polyclonal transgenic Neuro-2a lines; d) The transgene (HttQ15/HttQ138) copy number in the genomes of monoclonal (Q15m/Q138m) and polyclonal (Q15p/Q138p) Neuro-2a lines. The GAPDH-normalized Htt gene copy number in the (−) control line was referred to 1. e) 2^−ΔΔCt^ values for transgenic Neuro-2a lines with HttQ15 or HttQ138 expression and (f) Western blotting of lysates without/with doxycycline induction of HttQ15 or HttQ138 expression after 3 years of storage, cultivation and passaging. Total protein content in each sample was controlled by β-actin immunoreactivity. Data in Tables b) and f) and Fig. d) are represented as mean ± S.D. (n = 3).

Following a two-day incubation with doxycycline to induce transgene expression (+) or in the absence of induction (-), total RNA was extracted from each HttQ15 or HttQ138 candidate clone, from polyclonal cultures, and from control lines. Subsequently, the first-chain total cDNA was synthesized and then subjected to a quantitative PCR (qPCR) analysis using primers for exon 5 of the huntingtin gene. The selection of this exon was based on the optimal amplicon length for the qPCR method. The ΔΔCt method [45] was used to analyze results of the qPCR of Htt transgene expression (Fig. 1b). Doxycycline induction resulted in a pronounced overexpression of both HttQ15 and HttQ138 (Fig. 1b) in two polyclonal transgenic Neuro-2a lines with 45 ± 7-fold and 16 ± 4-fold increase in HttQ15 and HttQ138 expression, respectively. This is consistent with existing data that showing an increase of a similar order of magnitude in transgene expression compared to endogenous levels using the Sleeping Beauty system [46]. For further experiments, clones Q15#2 and Q138#3 were selected from four tested Q15 and Q138 monoclones (data not shown) due to their comparable transgene expression levels. HttQ15 and HttQ138 expression in selected monoclonal lines was about 15- and about 20-fold higher, respectively (Fig. 1b). The lengths of qPCR amplicons obtained for each of Neuro-2a monoclones were confirmed by agarose gel electrophoresis (Supplementary, Fig. S5), and a single 80 bp qPCR-product for each transgenic line was detected. The qPCR data shows successful transgene insertion in all poly- and monoclonal

Neuro-2a lines for both HttQ15 and HttQ138 genes. The levels of HttQ15 and HttQ138 protein production in poly- and monoclonal Neuro-2a transgenic lines were evaluated by Western blotting (Fig. 1c). The immunoblotting results are consistent with the expression levels data obtained by qPCR of both huntingtin variants in transgenic Neuro-2a lines. Integration numbers of transgenes in the genomes of all obtained transgenic cell lines were also estimated by qPCR (Fig. 1d). The Htt transgene copy numbers were calculated to be 5-10 integrated per cell population in monoclonal lines. This is in agreement with previous experiments using the Sleeping Beauty system [46, 47]. In Neuro-2a-HttQ15p/Q138p lines, the number of Htt gene integrates reaches 25-27, which explains the higher level of HttQ15/HttQ138 expression compared to monoclones.

Another significant criterion for the applicability and reliability of the Sleeping Beauty expression system integrated into Neuro-2a cells is the stability of the expression levels of HttQ15 or HttQ138, which persists over extended time periods. This is particularly relevant for cancer cell lines, including the Neuro-2a culture, since their genome instability, which accumulates with an increase in the number of passages, can negatively affect the expression level of the integrated transgene. Therefore, we re-evaluated the expression level of huntingtin types in our cell model (Fig. 1e, 1f) after 3 years of its use in our laboratory practice (including procedures for storage at −70°C, defrosting/freezing, cultivation and seeding of several dozen passages). Verification of HttQ15/Q138 mRNA expression levels showed that the main trends persisted for 3 years, and the levels of mRNA of huntingtin variants, according to RT-qPCR data (Fig. 1e), decreased mainly in polyclonal cells. The huntingtin protein production in Neuro-2a-HttQ15/Q138, however, has not undergone significant changes (Fig. 1f) compared to the state of 3 years ago, demonstrated stability of inducible HttQ15/HttQ138 protein expression.

### Transgenic Neuro-2a cells expressing mutant huntingtin demonstrated accumulation of the Htt aggregates and alteration in proteasome activity

The intracellular pattern of HttQ15 and HttQ138 expression in transgenic Neuro-2a lines (mono- and polyclonal cultures) was analyzed by immunohistochemical staining on the 7th or 14th day after the induction of transgene expression. As illustrated in Figure 2, the presence of Htt aggregates is clearly observable on the 14th day of HttQ138 expression.

**Figure 2.**
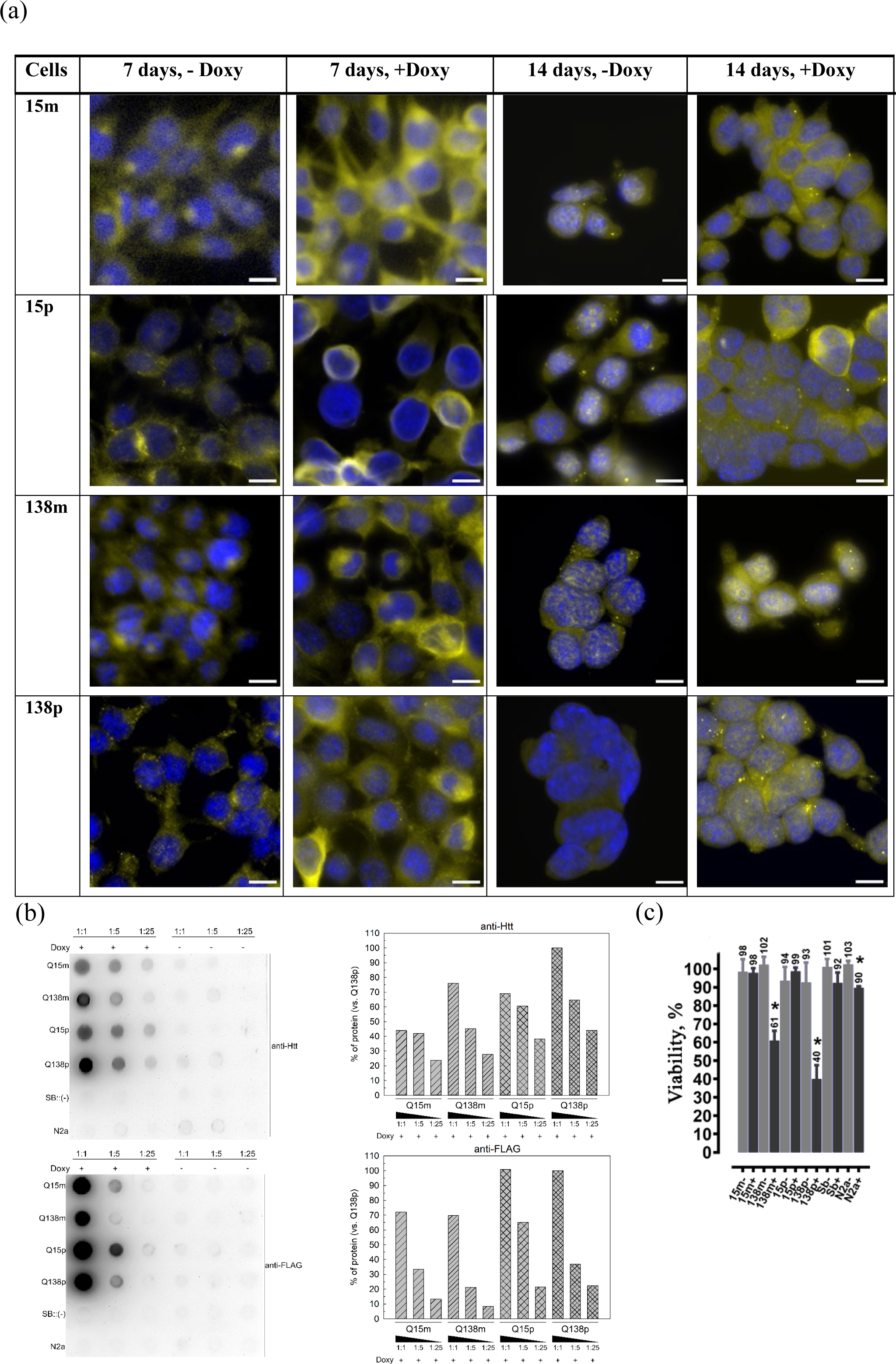
Generated transgenic cell line Neuro-2a/HttQ138 demonstrated accumulation of Htt aggregates after 14 days of HttQ138 overexpression, unlike cell line Neuro-2a/HttQ15. a) Immunofluorescence microscopy of transgenic Neuro-2a cells without (-)/with (+) overexpression of HttQ15 *or* HttQ138 after 7 and 14 days of doxycycline induction. Merge of yellow color which refers to huntingtin, and blue color (DAPI) – cell nuclei is presented. Scale bars are represented (10 μm). b) Filter trap assay of Neuro-2a transgenic cells’ lysates after 14 days of HttQ15/HttQ138 overexpression and without expression. Top: Htt (left panel) and FLAG-epitope (right panel) immunoreactivities are shown. Each sample was tested in 3 different dilutions (1:1, 1:5 and 1:25). Bottom: Relative intensities of HttQ15/HttQ138 (left panel) or FLAG-epitope (right panel) immunoreactivity in condition of transgenes’ expression after subtracting the corresponding signals without induction and normalization on total protein content in each sample (Ponceau S staining) reflect the specific presence of HttQ138 aggregates in cell lysates. Intensity of HttQ138p signal was referred to 100% in each experiment. c) Сell viability measured by MTT assay. \**p* < 0.05. Data in Fig. c) are represented as mean ± S.D. (n = 3).

After seven days of HttQ15/Q138 overexpression in our cell model, HttQ15 was found to be diffusely distributed almost throughout the cytoplasm and in the perinuclear region. Furthermore, immunoreactivity has been observed in neurofilaments for both huntingtin variants. Following a period of seven days, the presence of Htt-immunoreactive inclusion bodies was not detected. Prolonged expression for 14 days resulted in the formation of clearly distinguishable Htt-immunoreactive protein aggregates in both mono- and polyclonal Neuro-2a-HttQ138 lines (Fig. 2a; see also Supplementary, Fig. S6 for more details). Transgenic Neuro-2a-Q138p cells are characterized by a significantly larger number of mHtt aggregates compared to Q138m, which is evidently explained by a higher level of mutant huntingtin expression in some cells of the polyclonal line. Furthermore, it was observed that some HttQ138 inclusion bodies exhibited both intranuclear localization and cytoplasmic distribution (Fig. 2a). To confirm the presence of transgenic mutant huntingtin in the aggregates formed in the Neuro 2a-Q138m and Q138p lines after 14 days of expression, the lysates of each cell type were analyzed using the filter trap assay (Fig. 2b). Given the inability of large aggregates to traverse the membrane, visualization techniques such as Ponceau S staining and the use of antibodies directed against the N-terminal region of Htt or the C-terminal FLAG epitope can be employed to detect these aggregates. As demonstrated in Figure 2b, the lysates of Neuro-2a-HttQ138 transgenes exhibited the presence of aggregates containing mutant huntingtin, with the highest abundance of aggregated forms observed in Q138p cells. Neuro-2a-HttQ138m cells also demonstrate the presence of Htt-immunopositive aggregates, albeit to a lesser extent compared to Q138p, due to a lower Htt expression level in this transgenic line (Fig. 1b). Notably, even minimal amounts of aggregated Htt were detected in Q15m and Q15p cell lines, yet these aggregates were not observed in the control lines (SB and N2a) or in samples not treated with doxycycline. Upon sequential dilution of the samples (1:5, 1:25), a 10-20% decrease in aggregate content was observed. In the case of FLAG epitope immunoreactivity (Fig. 2b), no significant differences were observed between the cells expressing HttQ15 and HttQ138 (both in mono- and polyclonal lines). This can be explained by 1) a higher expression level of normal huntingtin compared to the mutant form in our model (Fig. 1b), 2) the unequal exposure of the FLAG epitope in Htt variants (probably FLAG epitope is more exposed in the HttQ15 structure and, accordingly, is available for antibody binding) and also by 3) cleavage of the C-terminus from huntingtin which leads to a lower content of FLAG-immunopositive signal in the aggregates.

As huntingtin aggregates has been demonstrated to disrupt intracellular proteolysis, the status of the proteasome system in Neuro-2a-HttQ15/Q138 monoclonal and polyclonal transgenic cell lines was evaluated after 14 days of induction. The results are presented in Fig. 3.

**Figure 3.**
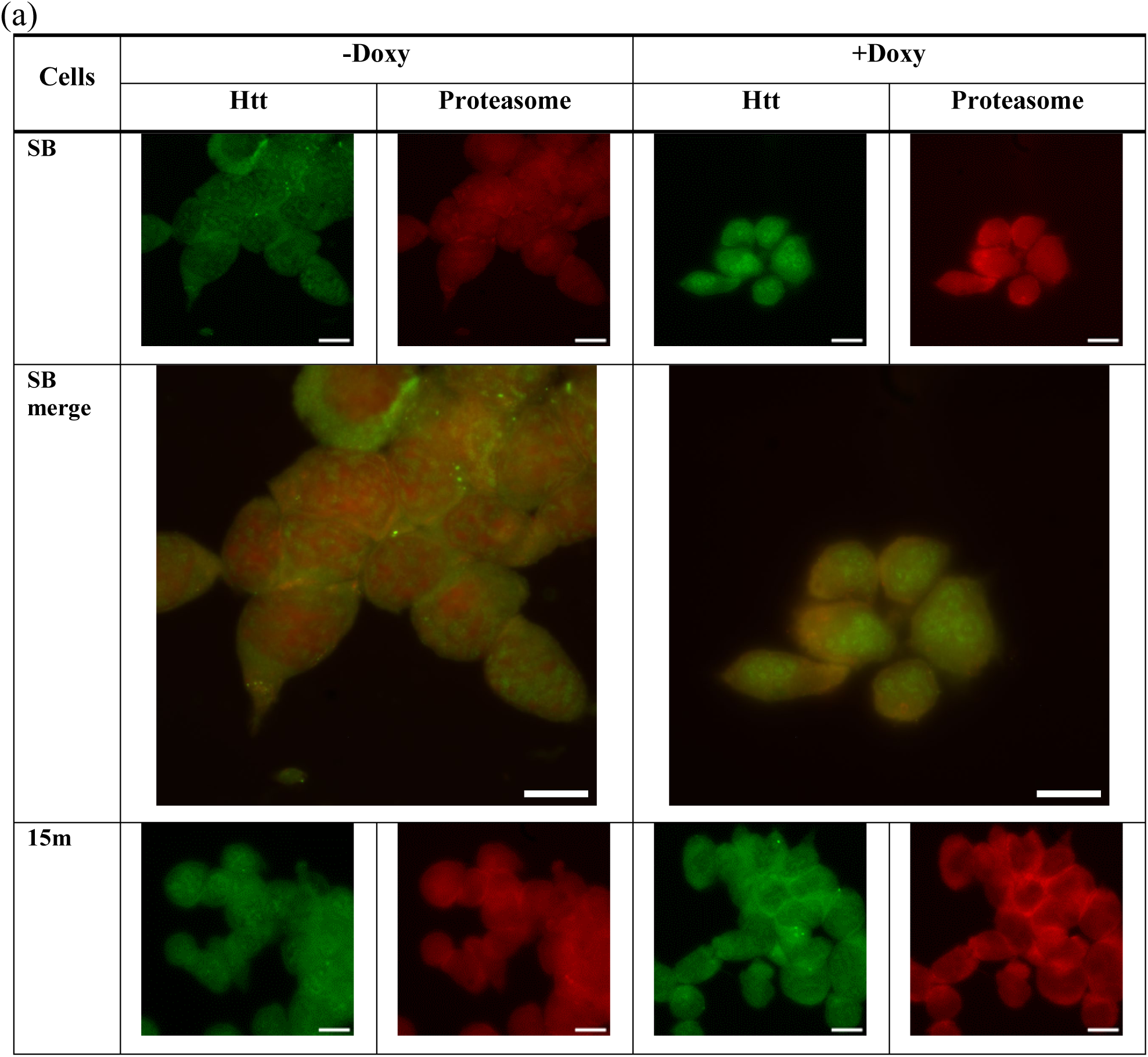
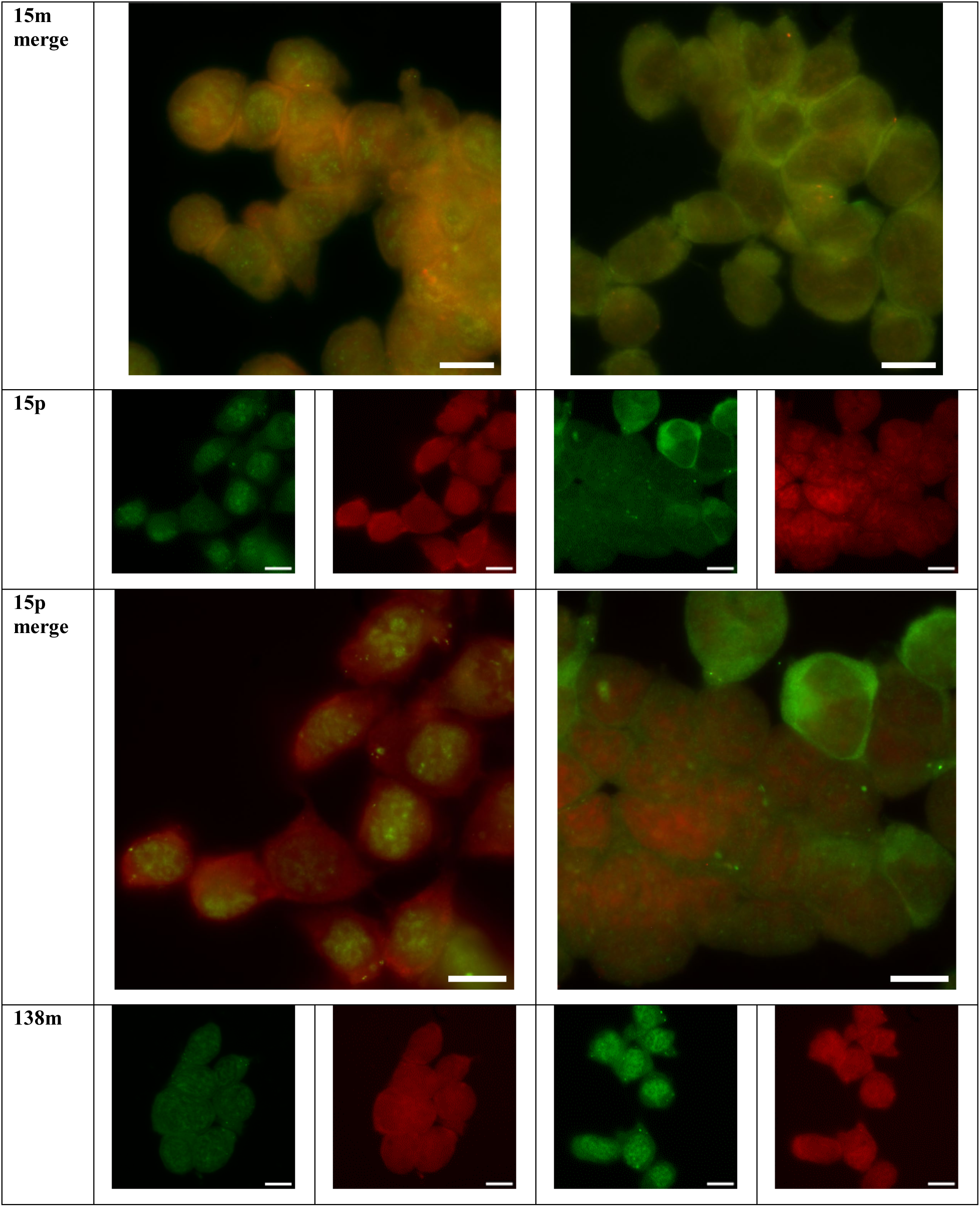
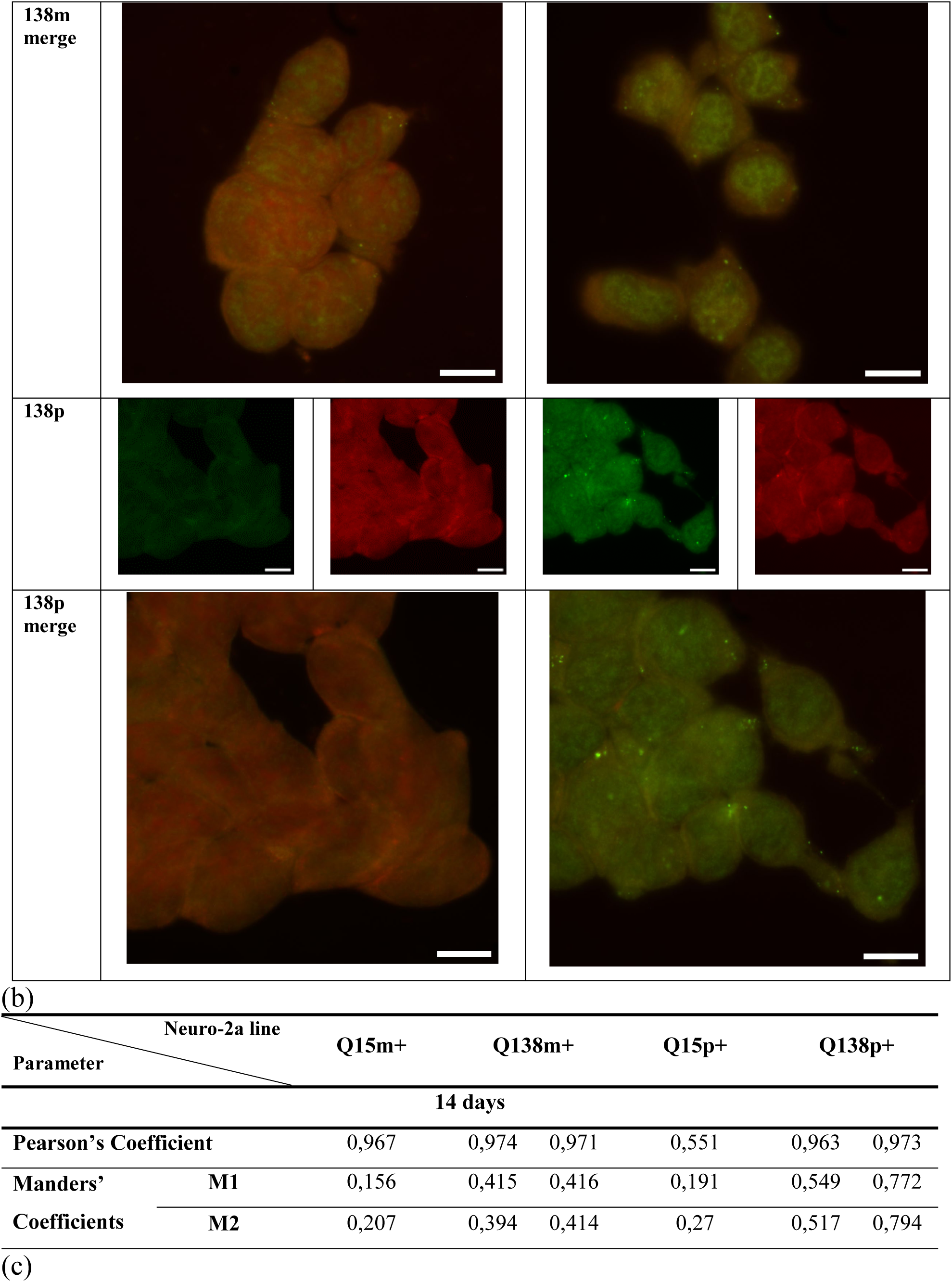
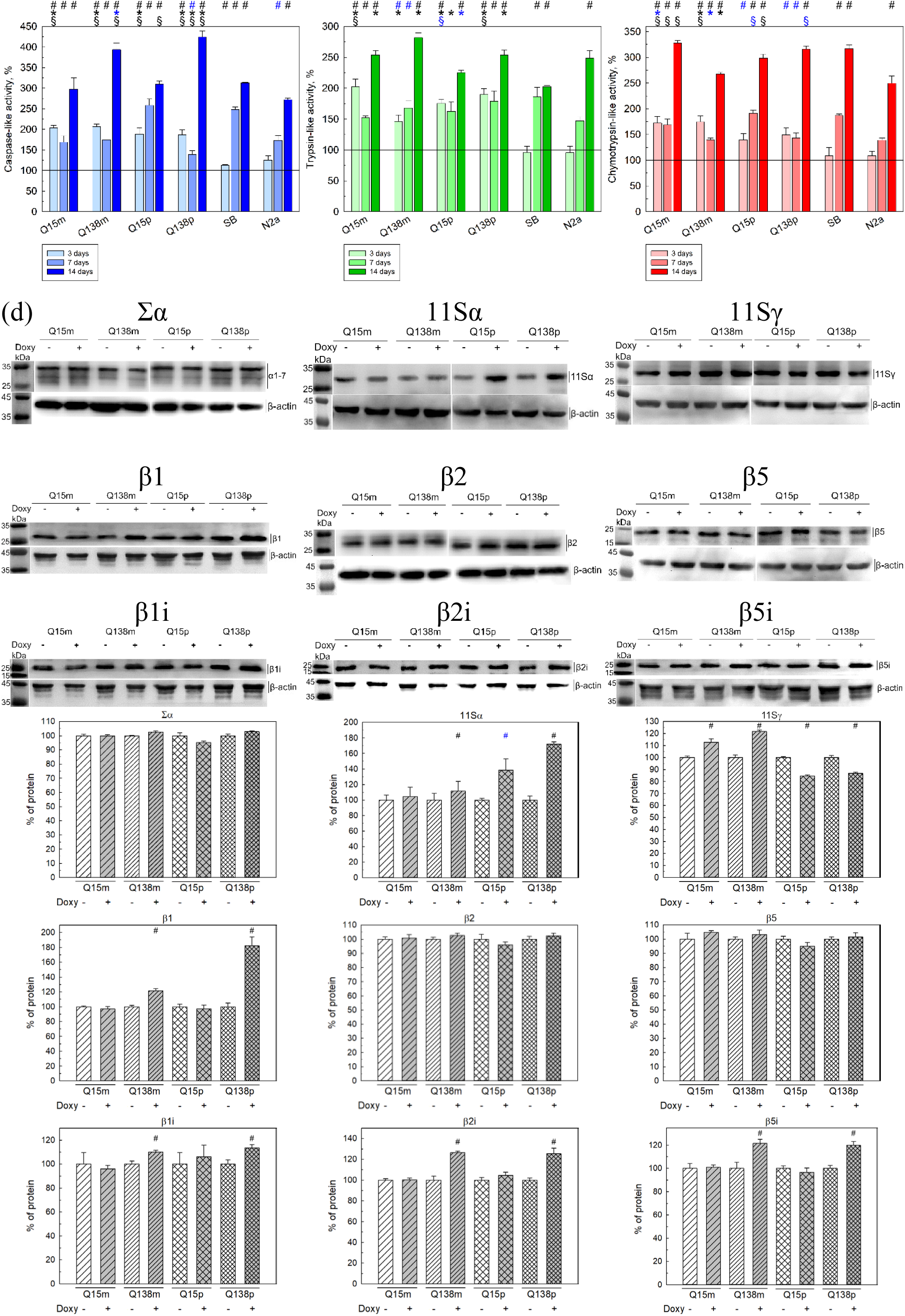
The proteasome is specifically activated in the Neuro-2a/HttQ138 transgenic cell line and shows colocalization with Htt aggregates/inclusion bodies. (a) Immunofluorescence microscopy of transgenic Neuro-2a/HttQ15/Q138 monoclonal (m) or polyclonal (p) cell lines without (-)/with (+) overexpression of HttQ15, HttQ138 after 7 and 14 days of doxycycline addition. Green fluorescence refers to huntingtin, red fluorescence refers to α1, 2, 3, 5, 6, 7 (marked as Σα) proteasome subunits. Scale bars are presented (10 μm). (b) Quantitative characteristics of huntingtin colocalization with the proteasome core subunits after 14 days of Htt expression in transgenic Neuro-2a-HttQ15/Q138m/p cells. (c) Three types of proteasome activity in transgenic Neuro-2a/HttQ15/Q138 cells during the overexpression of HttQ15 or HttQ138, as well as in control lines (SB, with transfection of an «empty» vector, and N2a, without transfection). Caspase-like (left panel), trypsin-like (middle panel) and chymotrypsin-like (right panel) proteasome activities were evaluated. The activity assay was conducted on days 3, 7 and 14 of induction. Proteasome activity in the doxycycline non-treated cell lines was referred to 100% in each experiment. d) Top panel: Western blotting of Neuro-2a transgenic cells lysates after 14 days of HttQ15/HttQ138 overexpression and without expression. All proteasomal α-subunits (α1, 2, 3, 5, 6, 7 or Σα), β1, β2, β5, immune variants of catalytical proteasomal subunits, 11Sα and 11Sγ regulators are visualized. Bottom panel: relative amounts of tested proteasomal subunits, 11Sα and 11Sγ proteins in transgenic Neuro-2a cells lysates after normalization on β-actin content. p, m – poly- and monoclonal cultures, respectively. Total protein content in each sample was controlled by β-actin immunoreactivity. Protein content in the doxycycline non-threated cells was referred to 100% in each experiment. ^#^*p* < 0,05, −Doxy vs +Doxy; \**p* < 0,05, +Doxy vs SB +Doxy; ^§^*p* < 0,05, +Doxy vs N2a +Doxy. Blue symbols indicate *p* < 0,1. Data in Fig. c) and d) are represented as mean ± S.D. (n = 3).

As indicated in Fig. 3a, HttQ138 but not HttQ15 is defined by a more localized accumulation of fluorescence signals. After 14 days of induction, distinct colocalization (40-80%) of huntingtin and 20S-proteasome was detected within the HttQ138-immunopositive inclusion bodies (Fig. 3a, merge, and Table on Fig. 3b). Quantitative characteristics of huntingtin colocalization with the proteasome after 14 days of expression in transgenic Neuro-2a lines demonstrate spatial convergence of proteasome core and HttQ138 in Htt-immunopositive aggregates.

To address the question of whether alterations in expression result in changes to proteasomal activity, three different peptide-based substrates were utilized, each corresponding to a specific type of activity. The measurements were conducted 3, 7 and 14 days after the induction of huntingtin expression, as well as in the absence of expression and in control Htt non-expressing lines (Fig. 3c). Following 3 days of HttQ15/HttQ138 expression, there was a clear activation (approximately 1.7-2.0-fold increase) of all types of proteasome activity in Neuro-2a monoclonal lines compared to doxycycline-untreated cells (Fig. 3c, light color). This effect is independent of the overexpressed form of huntingtin. It can be assumed that the 3-day duration of mutant huntingtin expression is insufficient for the manifestation of specific effects on these activity types. In both control lines, no significant changes in any type of proteasome activity were detected after treatment with doxycycline.

On the 7^th^ day of Htt expression (Fig. 3c, middle color), caspase-like proteasome activity remained virtually unchanged in Neuro-2a-HttQ15m and Q138m/p cells, with a slight increase observed in Q15p. In the control Neuro-2a cells, a slight activation was detected, reaching approximately the same level of activation as in Q15m and Q138m/p cells. An examination of trypsin-like activity revealed no significant alterations in the Neuro-2a-HttQ15p and Neuro-2a-HttQ138m/p lines compared to the 3^rd^ day (Fig. 3c, middle panel). However, a modest decrease in trypsin-like activity was observed in Neuro-2a-HttQ15m line. Chymotrypsin-like activity exhibited minimal alterations in the Neuro-2a-HttQ15m and Neuro-2a-HttQ138m/p on (Fig. 3c, right panel) lines compared to the 3rd day, with a modest increase observed in Neuro-2a-HttQ15p. This phenomenon is presumably attributable to the establishment of steady-state level of these proteasome activity types in response to the expression level of normal huntingtin. Following a week of treatment with doxycycline, a significant increase in all three types of the proteasome activity was observed in the control Htt-non-expressing line, particularly in the SB line, when compared to the 3^rd^ day of treatment. This increase was likely due to significant overexpression of luciferase in response to the addition of doxycycline. Consequently, the cells attempt to degrade excessive protein.

Prolonged treatment with doxycycline during 14 days (Fig. 3c, dark color) resulted in a significant elevation of chymotrypsin-like proteasome activity (2.7-3-fold compared to doxycycline-untreated cells) in all transgenic Neuro-2a lines. However, this effect was comparable to doxycycline activation in control SB and Neuro-2a lines. A similar, but less pronounced effect (Fig. 3c, 2.3-2.8-fold increase) was observed for trypsin-like proteasome activity (Fig. 3c, middle). This particular activity type, however, has been observed to increase specifically in response to excessive production of HttQ138 (by approximately 30-35% compared to the Neuro-2a-HttQ15m/p lines). Of particular note is the marked increase (3-4-fold in comparison to cells without induction) in the caspase-like proteasome activity, as illustrated in (Figure 3c, left). In this case, the HttQ138 overexpression effects substantially exceed the nonspecific doxycycline-induced proteasomal activation. It is worth noting that all the observed changes in proteasomal activity types are primarily attributable to the proteasome contribution, since no significant changes in the cellular background protease activity (in the presence of the MG132 proteasome inhibitor) were detected during the two-week experiment (Supplementary, Fig. S7).

After 14 days of HttQ15/HttQ138 expression in Neuro-2a transgenic lines, no substantial alterations were observed in either the total content of the 20S proteasome (Fig. 3d, bottom panel) or the expression level of the β2 and β5 subunits (the increase in the β2 or β5 content upon HttQ15/HttQ138 overexpression is not more than 10%). However, it was established that mutant huntingtin expression leads to an increase in the level of β1 subunit by 20% in the monoclonal and 70% in the polyclonal Neuro-2a lines (Fig. 3d, bottom panel). Additionally, after 14 days of mHtt overexpression the content of each of the immune proteasomal subunits increased by 10-25% in both HttQ138m/p lines, but not in HttQ15m/p cell lines (Fig. 3d, bottom panel). It is also possible that the proteasomal response to HttQ15/HttQ138 overexpression may be simultaneously mediated by the action of proteasome regulators. Indeed, a marked increase in the 11Sαβ-regulator amount (by approximately 60%) is observed upon both HttQ15 and HttQ138 overexpression (Fig. 3d, top panel). Therefore, the activity measurements demonstrated that doxycycline treatment caused total proteasome activation, with a specific and pronounced elevation in caspase-like activity observed in transgenic Neuro-2a-HttQ138m/p cell lines. The specific activation in transgenic Neuro-2a-HttQ138m/p cells was clearly associated with elevated expression of the β1 catalytic subunit, whereas total proteasome activation in these cells – with stimulation of immune proteasome subunits expression. In parallel, it was originated from elevated expression of the 11Sαβ activator (Fig. 3d).

### Transgenic Neuro-2a_SB_HttQ15/Q138 cells revealed variation in autophagy

It is well established that Htt aggregates are the substrates for degradation by autophagy [48]. In this context, an additional evaluation was conducted on the effect of HttQ15/HttQ138 expression on the level of LC3B, an essential marker of the autophagy process, as well as on the amount of lysosomal proteinase cathepsin D, cathepsins B and D activity. The results obtained are summarized in Figure 4.

**Figure 4.**
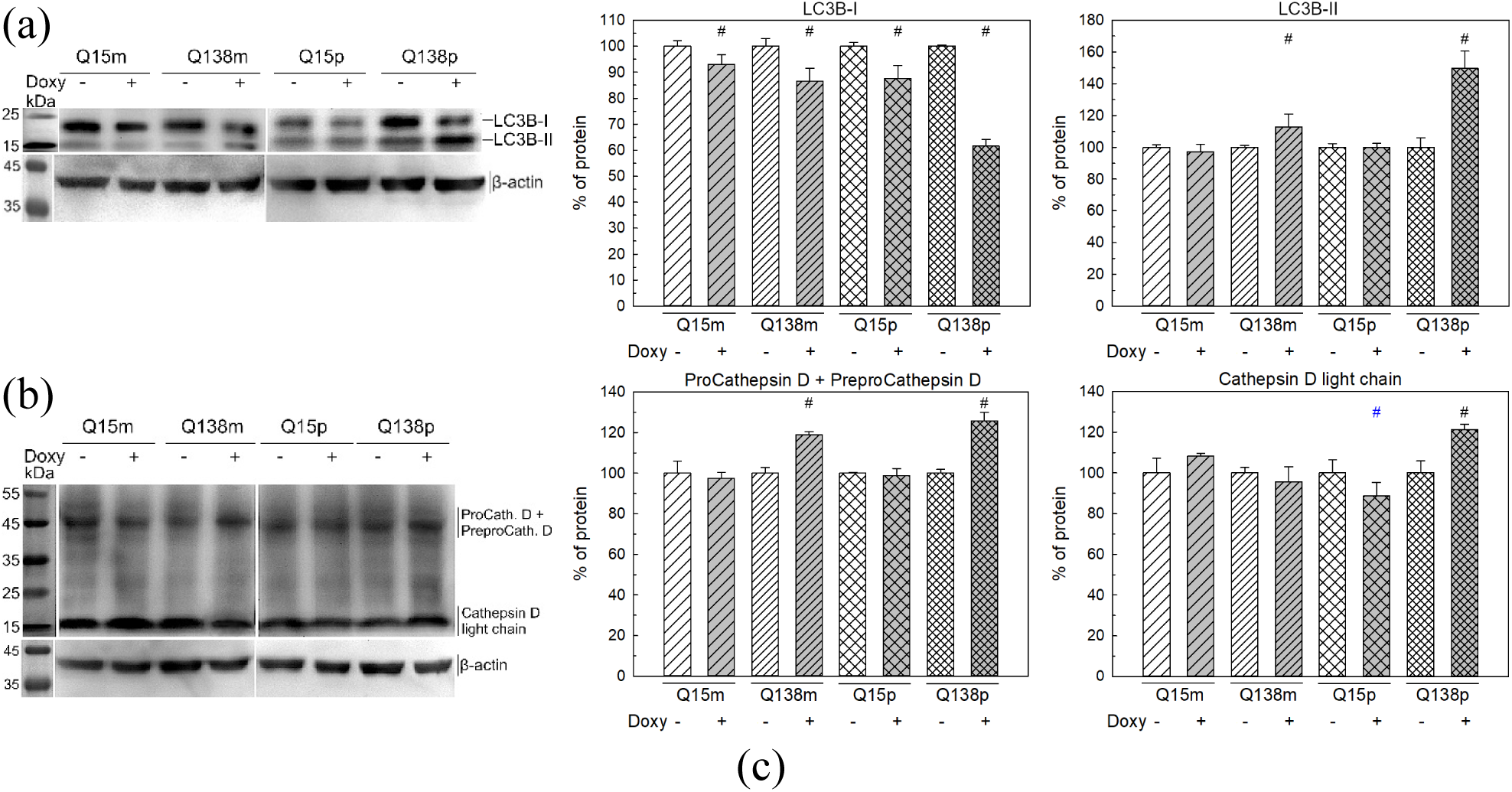
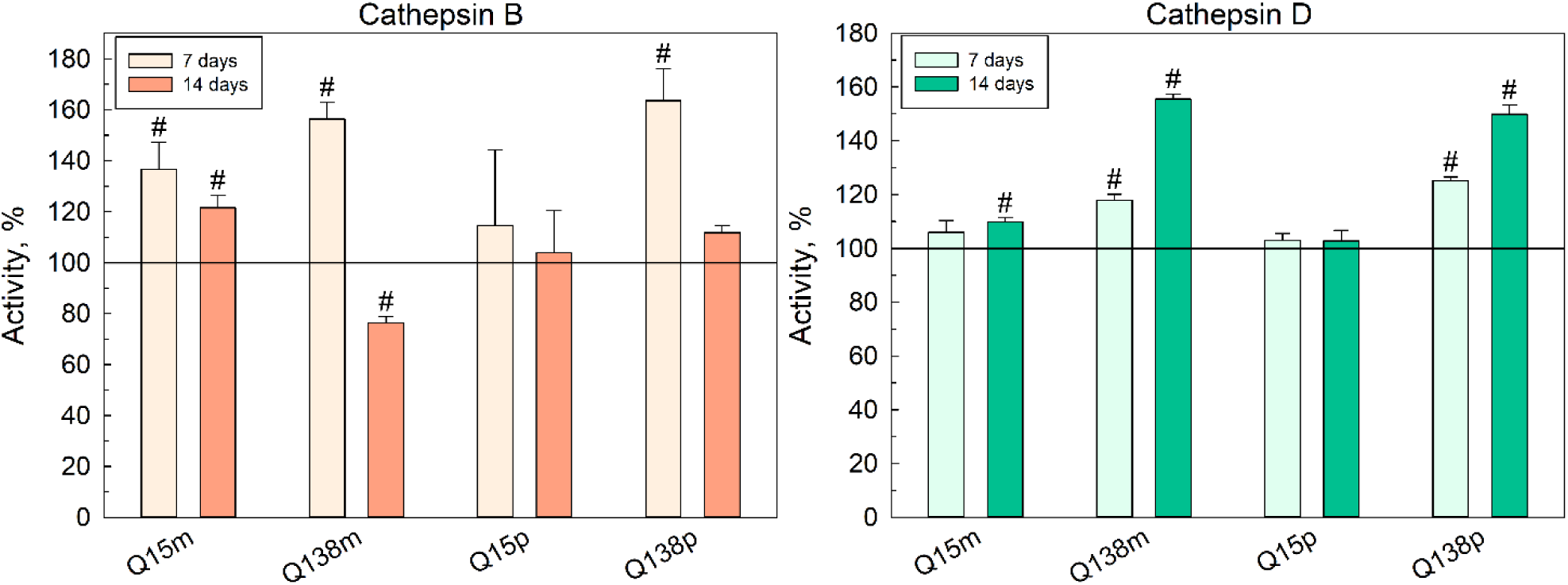
Activation of autophagy is linked to mutant Htt overexpression. Left panel: Western blotting of Neuro-2a transgenic cells lysates after 14 days of HttQ15/HttQ138 overexpression and without expression. Right panel: relative amounts of tested proteins in transgenic Neuro-2a cells lysates after normalization on β-actin content. a) LC3B-I/II, b) cathepsin D proteins are visualized. Protein content in the doxycycline non-threated cells was referred to 100% in each experiment. ^#^*p* < 0,05, −Doxy vs +Doxy. Blue symbols indicate *p* < 0,1. c) Intracellular activity of cathepsin B using Z-FR-AMC substrate (left) and cathepsin D using Abz-A_2_P_2_-DeD substrate (right) in transgenic Neuro-2a cells during the overexpression of HttQ15 or HttQ138. Pale color indicates 7 days and bright color indicates 14 days of doxycycline induction. p, m – poly- and monoclonal cultures, respectively. Data are represented as mean ± S.D. (n = 3).

Only minor changes could be detected in the LC3B-I/II levels upon HttQ15/HttQ138 overexpression in monoclonal lines and in Q15p line (Fig. 4a, right). Western blotting results demonstrated the decrease in LC3B-I expression level of no more than 20% in monoclones, as well as in Q15p line, and more pronounced decrease (about 40%) in Q138p cells. This decline in the LC3B-I level with a simultaneous increase in the amount of LC3B-II was observed in Neuro-2a-HttQ138m/p cell lines demonstrating an intensification of LC3B lipidation, which is particularly pronounced in the Q138p line (where the level of LC3B-II increases by approximately 50% in comparison with the absence of mHtt expression). It is also worthy of note that HttQ138 overexpression in the Neuro-2a polyclonal line resulted in the increase of cathepsin D level by approximately 30% (Fig. 4b, right). The elevation of lysosomal cathepsin B activity (Fig. 4с, left) was primarily observed following 7 days of expression induction. However, after 14 days, the increased activity was retained exclusively in the Neuro-2a/HttQ15m lines. In general, lysosomal cathepsin B activity was found to be normalized in the Neuro-2a/HttQ15/HttQ138p lines and decreased in the Neuro-2a/HttQ138m cells. On the other hand, the activity of another lysosomal cathepsin D (Fig. 4c, right), demonstrating an increase in Q138m/p lines by 20-30% already on 7^th^ day of huntingtin overexpression, maintains and enhances this trend after 14 days of induction, reaching an elevation of 50-60% in transgenic lines with mutant huntingtin production, which corresponds to an augmented cathepsin D expression in Q138m/p lines according to the results of Western blotting (Fig. 4b, right).

## Discussion

This study discloses the proteostasis alteration in a mutant huntingtin-overexpressing cell lines that have been derived from the neuronal line Neuro-2a. The Neuro-2a cell line is a frequently employed mouse neuroblastoma that is characterized by its stability, relatively rapid growth rate, and ease of handling. The generation of Neuro-2a-HttQ15/Q138 cell lines was achieved through the application of the Sleeping Beauty system, that facilitates the inducible expression of the *HTT* gene using doxycycline. The qPCR results obtained have indicated that a sufficient number of normal or mutant huntingtin gene copies were integrated into the genome of Neuro-2a cells, despite the large size of the Htt gene. It is important to note that this is the first time that such a genomic feature has been evaluated for full-length huntingtin variants in similar cellular expression systems. Two variants of the Neuro-2a-HttQ15/Q138 cell line were created – a polyclonal (p) and a monoclonal (m). It is important to acknowledge the inherent heterogeneity of the Neuro-2a-HttQ15p/Q138p lines, which comprise cells exhibiting different levels of transgene expression. It was calculated that Htt transgene copy numbers in monoclonal lines were substantially lower than in polyclones, where it reaches 25-27 (Fig. 1d). It is noteworthy that the number of integrations could be further reduced by reducing the amount of transposon DNA during transfection [49]. The Western blotting results confirmed high levels of HttQ15 and HttQ138 protein production in poly- and monoclonal Neuro-2a lines (Fig. 1c), and these results are consistent with the expression levels data obtained by qPCR of both huntingtin variants in transgenic Neuro-2a lines. Thus, following the validation of the HttQ15 and HttQ138 transgene expression levels in Neuro-2a cells using two independent approaches, it can be concluded that the Sleeping Beauty system is applicable for stable transfection of large proteins in the selected cell line. The Neuro-2a-HttQ15/Q138 transgenic cell lines exhibited relatively stable HttQ15/Q138 protein expression characteristic (Fig. 1e, f). Consequently, we also demonstrated for the first time that the created transgene Neuro2a cell line with expression of huntingtin variants maintains the stability of its functioning for a period of at least three years, which allows for the exploitation of this HD model for long-term experiments and observations.

The hallmark of HD at the cellular level is the presence of Htt-immunoreactive inclusion bodies [50, 51]. The process of huntingtin expression in Neuro-2a cells was found to result in gradual Htt accumulation, with the protein becoming diffusely distributed throughout the cell, except of the nucleus after a 7-day period of HttQ15/HttQ138 expression. The prolonged 14 days-expression resulted in the formation of clearly discernible Htt-positive aggregate in Neuro-2a-HttQ138m/p cell lines. Neuro-2a-HttQ138p cells demonstrates higher ability to accumulate mHtt aggregates compared to Neuro-2a-HttQ138m, and part of HttQ138 aggregates localize not only in the cytoplasm but also within the nucleus. The presence of mHtt nuclear aggregates is also an essential distinguishing feature of the HD molecular pathogenesis. According to some reports, the nuclear aggregates formation occurs at an early stage of the disease [52] and entails the most serious toxic effects for neurons, since it leads to global violations in many aspects of the neuronal functioning, for example, to a transcription dysregulation of genes responsible for maintaining proteostasis, the production of neurotrophic factors, etc [53]. The filter trap assay data indicates the presence of HttQ138 in aggregates within the Neuro-2a transgenes lysates with Neuro-2a-Q138p cells having the highest number of aggregated forms which is consistent with the immunofluorescence microscopy results. Filter trap assay with FLAG epitope detection did not show a significant difference between the cells expressing HttQ15 and HttQ138, rather more difference was observed between mono- and polyclonal lines. One potential explanation is the elevated intracellular proteolysis during Htt overexpression. It was demonstrated that under conditions of transient transfection in HEK293T cells, active intracellular proteolysis occurs for both normal and (to a greater extent) mutant forms of Htt 1st exon (httN233) [54]. In the cellular model under investigation, the cleavage of C-terminal Htt fragments, which contain the FLAG epitope [40], is possible, and probably worsens its visualization. Thus, using the filter trap assay, we were able to verify that the inclusion bodies we observed in model Neuro-2a cell lines contain transgenic huntingtin with FLAG epitope. Neuro-2a-based cellular model developed in this study has been demonstrates the most characteristic distinctive HD molecular features related to different stages of the disease.

Numerous data indicate that intracellular protein aggregates in various HD models actually contains the proteasome complex components (20S-core particle and others) [9, 55]. The effect of HttQ15/HttQ138 expression during 3 days was most noticeable on caspase-like and in less extent on trypsin-like proteasome activity in all transgenic cells, and these results are consistent with our data on the transient expression (during 72 h) of huntingtin in HEK293T cells, where an increase in caspase-like proteasome activity was most prominent [40]. Significant proteasome activation in control Htt-non-expressing lines (especially in SB) was detected after 7 days of doxycycline treatment. It was shown in a mouse model that oral administration of 1,5 mg/ml doxycycline for 30 days leads to no more than 10% increase in the chymotrypsin-like proteasome activity [56]. However, in our cellular model, 1,5-2,0-fold increase in each proteasome activity type was observed in response to doxycycline treatment. It is also possible that a significant change in activity occurs under the accumulation of the protein (luciferase) that was encoded in the «empty» SB vector. The raw data (Table S1) depicts a rather low chymotrypsin- and caspase-like activity in these cells without doxycycline. Consequently, after 7 days of doxycycline supplementation, alterations in proteasome activity can be attributed to the combined impact of doxycycline addition and Htt overexpression, making it impossible to discriminate between them.

Following 14 days of doxycycline supplementation, the most significant alteration was observed in caspase-like proteasome activity, and the HttQ138 expression led to the most substantial change in activity (Fig 3c, left). The proteasome catalytic subunit β1, which possesses caspase-like activity, has been demonstrated to cleave peptide bond after negatively charged amino acid residues (commonly, aspartate and glutamate) and comprise the most hydrophilic S1 specificity pocket [57, 58], therefore this subunit may be capable of hydrolyzing polyQ fragments. It was recently determined [59] that *in vitro* hydrolysis of a synthetic polyglutamine Q_10_ substrate involves predominantly caspase-like proteasome activity, while for a shorter Q_5_ substrate hydrolysis the β1 and β2 subunits could contribute together. A global increase in the caspase-like proteasome activity under HttQ15/HttQ138 overexpression may also be connected with the involvement of proteasome regulators, such as 19S or 11S proteins. It has been established that the 11Sαβ regulator enhances the rate of hydrolysis of fluorogenic peptide substrates by the 20S proteasome for all three substrate specificity types [60, 61]. Similarly, the purified eukaryotic 20S/26S proteasome has been observed to cleave only the flanking sequence of synthetic peptide Q_10_ substrates or perform hydrolysis after the first glutamine residue, but the proteolysis rate increases significantly in the presence of the 11Sαβ regulator [59, 62]. As demonstrated in human HEK293 Tet-Off cells, the insoluble HttQ83 exon 1 cytoplasmic aggregates colocalized with 11S regulator [25, 63, 64]. The cytoplasmic localization of 11Sαβ expression, together with the evidence of its colocalization with mHtt aggregates [65, 66], supports the hypothesis that 11Sαβ activator could be involved in the specific regulation of proteasome activity during the proteolysis of mHtt. Consequently, the observed effects of HttQ15/HttQ138 overexpression on various types of proteasome activity may be a combination of mHtt toxic effects on the UPS and the compensating action of proteasome regulators. In order to gain a more complete understanding of how regulator proteins impact mHtt proteolysis by proteasome in this Neuro-2a based HD cellular model, further studies are required, including testing of different proteasome activities under conditions of repression/overexpression of 19S and/or 11S.

In order to ascertain whether the overexpression of HttQ15/HttQ138 influences the expression levels of proteasomal core α- and β-subunits as well as regulatory proteins, their quantity was estimated by Western blotting of Neuro-2a lysates following 14 days of HttQ15/HttQ138 expression in comparison to the absence of induction. No significant (more than 10%) changes were detected in either the total content of the 20S proteasome or the expression level of the β2 and β5 subunits. However, a marked increase in the β1 subunit level (by 50-60%) was detected specifically under the influence of mHtt toxicity and pronounced accumulation of 11Sαβ-regulator (by 60%) was observed upon both HttQ15 and HttQ138 overexpression. Certainly, the pronounced increase in the β1 subunit production, compared with other proteasome catalytic subunits in Neuro-2a-Q138m/p cells, is indicative of the most significant and selective enhancement of the proteasomal caspase-like activity after 14 days of huntingtin variants overexpression (Fig. 3c, left). The level of immune proteasomal subunits increased by 10-25% in HttQ138m/p lines, reflecting activation of the immune proteasome, which is more active than the constitutive proteasome. The 11Sαβ regulator, which forms complex predominantly with 20S immune proteasome, has been demonstrated to play a significant role in the pathogenesis of polyQ diseases, with the capacity to activate the proteasome for the degradation of polyQ-containing proteins [10]. Hence, an increase in the expression level of the 11Sαβ regulator may serve as a cellular response to proteotoxic stress caused by excessive accumulation of mutant huntingtin. The 11Sγ protein level demonstrates a comparable tendency to rise slightly in Neuro-2a-HttQ15/HttQ138m, yet in polyclonal lines there is a modest decline in 11Sγ expression, which correlates with proteasome activity. It has been established that 11Sγ specifically enhances the trypsin-like proteasome activity, partially inactivating the other two types [60, 67]. However, after 14 days of huntingtin expression, the maximal increase in proteasome activity was observed for chymotrypsin-like and caspase-like types (Fig. 3c, left and middle). According to recent data, increased expression of proteasomal immunosubunits could provide a generalized neuroprotective mechanism in various models of neurodegenerative diseases [68–70]. As reported in [71], the levels of LMP2 (β1i) and LMP7 (β5i) proteasome immunosubunits were found to be elevated in the HD94 mouse brain homogenate. LMP2 immunosubunit is also upregulated in other neurodegenerative diseases, in particular this effect has been observed in mouse models of Alzheimer’s disease [72]. A recent study has established a link between β5i upregulation and the development of Parkinson’s disease in patients [73]. Concurrently, the induction of the proteasome immune subunits, can be caused not only by IFN-γ, but also by the 11S regulator, as has been demonstrated previously [74, 75]. Presumably, this particular phenomenon is manifested in the presented HD model, where the 11Sαβ overregulation provokes a slight (by 20-25%) but significant increase in the levels of all three types of immune subunits (Fig. 3d, bottom panel). At the same time, a decrease in the expression of 11Sγ may correlate with the activation of certain immune subunits, for example, LMP7 [76], which is also reflected in the features of proteostasis in our cellular HD model.

It is widely acknowledged that huntingtin typically functions as a critical component in the process of the autophagosome assembly. The expression of mutant huntingtin, itself being an autophagy substrate, has been shown to pathologically exacerbate the accumulation of either empty autophagy vesicles or autophagy vesicles containing unutilised cargo [48, 77]. The finding that mHtt functions as a substrate for lysosomal proteases, including cathepsins, further substantiates the involvement of the autosomal-lysosomal pathway in mHtt proteostasis [18, 78]. Only minor changes in the LC3B levels were detected upon HttQ15m/HttQ138m overexpression, as well as in Q15p line (Fig. 4a). Presumably, at lower levels of HttQ15/HttQ138 expression in monoclonal lines, an overregulation of autophagy can still be observed, whereas mainly in in Q138p line an acute accumulation of HttQ15/HttQ138 leads to an overload of the autophagosomal-lysosomal system. The overexpression of HttQ15/HttQ138 has been demonstrated to affect the levels of both the initial form of LC3B (LC3B-I), and the processed form LC3B-II, which is post-translationally modified with a phosphatidylethanolamine residue [79]. This modification allows LC3B to participate in the processes of autophagosome membrane expansion and their fusion with lysosomes [80]. Therefore, monitoring the content of LC3B-II provides information about the intensity of autophagy in created HD cellular model. The expression of HttQ138 leads to an intensification of LC3B lipidation, a process which is especially pronounced in the Q138p line. Presumably, at lower levels of mHtt expression in the Neuro-2a monoclonal line, the utilization of toxic forms of mHtt remains to a certain extent under the control of the UPS, and does not require significant activation of the autophagic flux. In the Neuro-2a-HttQ138p line, the presence of a high concentration of mHtt has been shown to induce a malfunction in the proteasome, resulting in compensatory overregulation of autophagy. This phenomenon has been observed in numerous analogous model systems and also patients with HD [48, 81]. Noteworthy, HttQ138 overexpression in the Neuro-2a polyclonal line resulted in the increase in cathepsin D level by approximately 30% (Fig. 4b), and also leads to a corresponding increase in enzyme activity by almost 60% (Fig. 4с, right). This lysosomal enzyme is a major aspartyl endopeptidase in the mammalian brain, and the presence of cathepsin D in neurons has been previously been demonstrated [82]. Consequently, not only an increase in the formation of autophagosomes has been observed, but also an increase in the activity of lysosomal enzymes. Increased expression of cathepsins B and D has been shown to effectively reduce both full-sized mHtt and its N-terminal fragment levels in transfected HEK293 cells. Neuroprotective effects were also observed upon the individual expression of cathepsins B or D in primary neurons [18, 78]. Hence, elevated cathepsin D expression and activation can also be a compensatory mechanism that activates in response to UPS overload caused by mHtt expression. From other side, lysosomal cathepsin B activity was increased following 7 days of HttQ15/HttQ138 expression induction, but returned to baseline levels after 14 days in all transgenic lines, with the exception of Neuro-2a-HttQ15m. It is important to note that no significant changes in the expression levels of the subunits/proteins under discussion were observed after 3 days of Htt/mHtt expression (Fig. S8). To summarize, the mutant huntingtin overexpression in the HD cellular model developed in this study has been shown to trigger cellular responses that appear to be aimed at reducing the mHtt toxicity and rebalancing the proteostatic network.

## Conclusion

In this study we have established a new transgenic HD cellular model with inducible expression of full-length normal and mutant Htt forms, starting with the construction of vector plasmids and proceeding with the description of the effects of HttQ15/HttQ138 overexpression on the functioning of the proteasome complex and other participants of the cellular proteostasis system. Furthermore, in the constructed model, we were succeeded to detect the most striking molecular hallmarks of HD pathology (the formation of intracytoplasmic and intranuclear mHtt aggregates and their colocalization with the 20S proteasome). The findings indicate that the expression of HttQ138 led to a substantial alteration in caspase-like proteasome activity, with trypsin-like activity also being enhanced, albeit to a lesser extent. Simultaneously, the expression of both full-size HttQ15 and HttQ138 resulted in significant increase in the 11Sα-regulator level and a concurrent decrease in the level of 11Sγ-regulator, particularly in the Neuro-2a-HttQ138p line. Furthermore, the expression of HttQ138 has been demonstrated to intensify LC3B lipidation and to considerable increase in lysosomal cathepsin D expression and activity, resulting in an increase in functional autophagosome formation. Taken together, these results reflect alterations in the proteostasis network upon HttQ138 expression and demonstrate mHtt proteotoxicity. Further elucidation of the qualitative/quantitative characteristics of this Neuro-2a cellular model (i.e. the expression pattern UPS components) may help to reveal previously unknown features of this complex proteinopathy. This is of significant importance for the advancement of understanding the pathological mechanisms that occur during the manifestation and development of HD pathogenesis. Furthermore, the results of this study will facilitate the development of new treatment strategies and the identification of potential therapeutic targets.

## Acknowledgments

The study was supported by the Russian Science Foundation grant number 25-24-00260. Authors are grateful to M.V. Lomonosov Moscow State University Program of Development for the opportunity to use Olympus FV1000/IX81 for cell imaging.

## Author Contributions

Conceptualization, A.V.B.; methodology, N.N.G., T.V.B., A.A.V., M.P.R., M.I.Z. and A.A.E.; validation, all authors; formal analysis, A.V.B., A.A.V. and N.N.G.; investigation, N.N.G., A.A.V., A.A.E., and A.V.B.; resources, A.V.B., M.P.R., M.I.Z. and A.A.E.; writing—original draft preparation, A.V.B. and N.N.G.; writing—review and editing, all authors.; visualization, A.V.B., N.N.G., A.A.V. and A.A.E.; supervision, A.V.B.; funding acquisition, A.V.B. All authors have read and agreed to the published version of the manuscript.

## Supplementary Materials

**Fig. S1.**
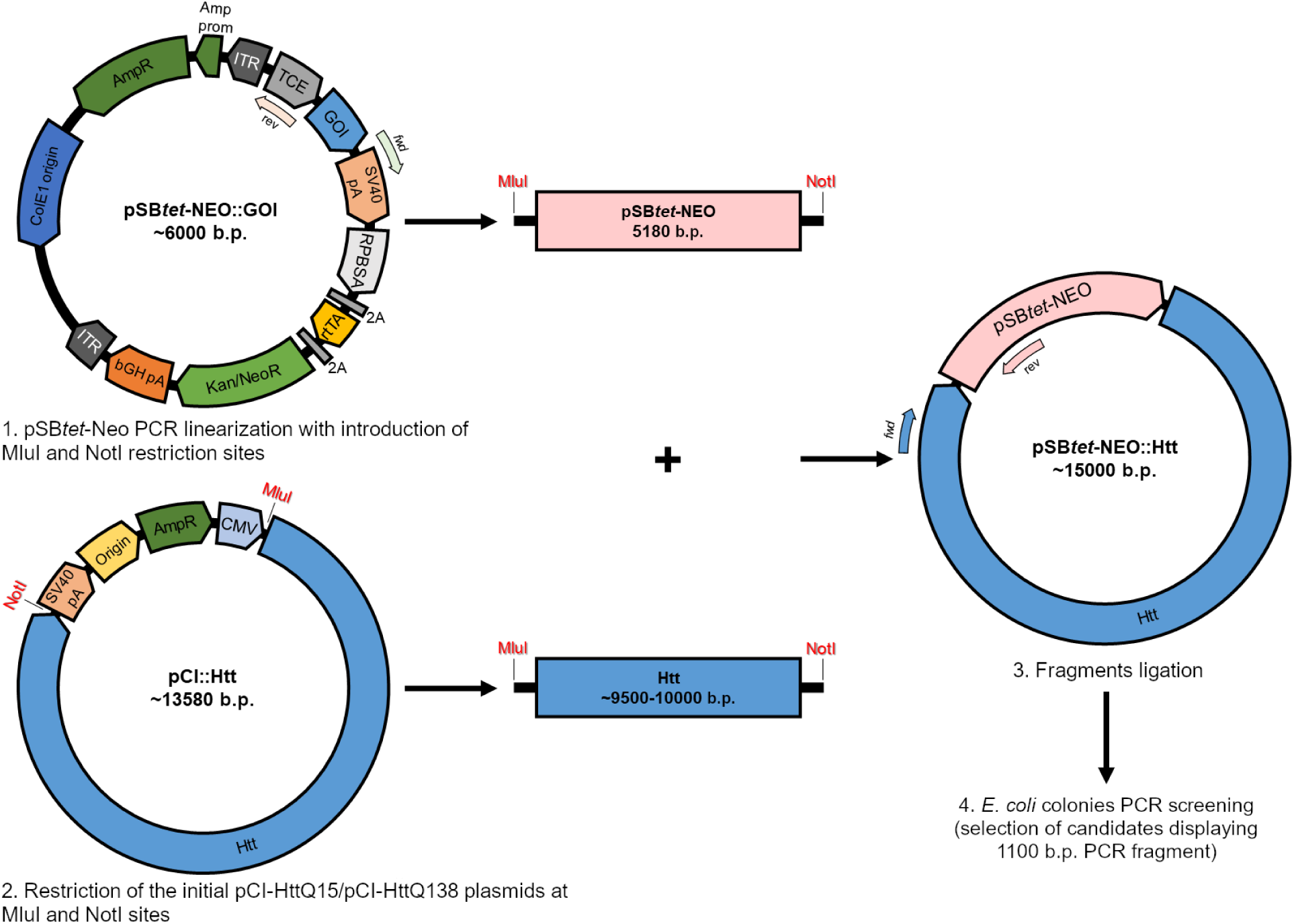
Scheme of Htt genes cloning into the pSB*tet*-Neo expression vector. ITR — inverse terminal repeats, TCE — tetracycline-inducible promoter, GOI — studied gene, RPBSA — synthetic constitutive promoter, 2A — flanking fragments with proteolysis sites, rtTA — tetracycline-inducible transactivator, Kan/NeoR, AmpR — kanamycin/neomycin and ampicillin resistance genes, respectively, SV40 pA and bGH pA — polyadenylation signals, CMV — CMV immediately-early enhancer/promoter, Htt — normal or mutant huntingtin gene.

**Fig. S2.**
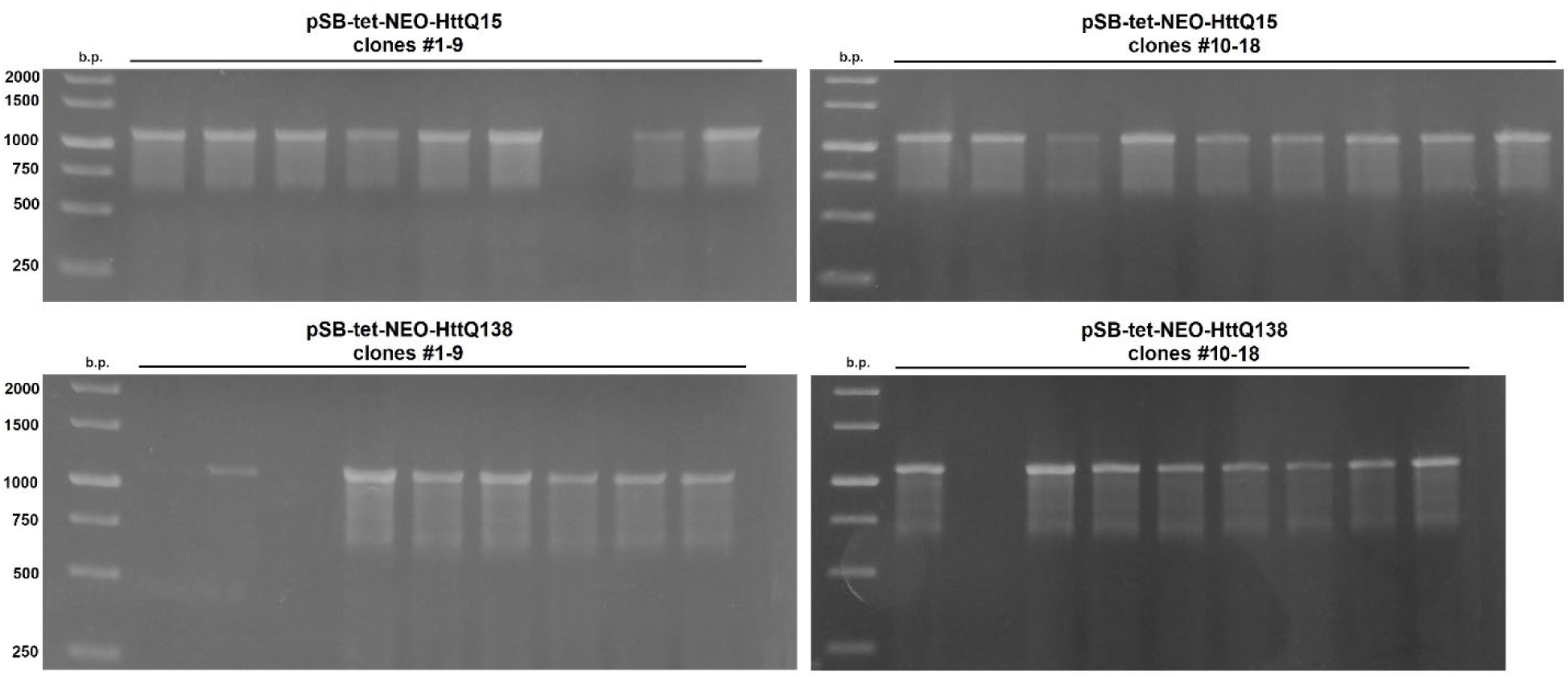
Agarose electrophoresis of PCR products obtained during the PCR selection of *E. coli* X1-Blue clones containing the pSB*tet*-Neo-HttQ15 or pSB*tet*-Neo-HttQ138 constructs.

**Figure S3.**
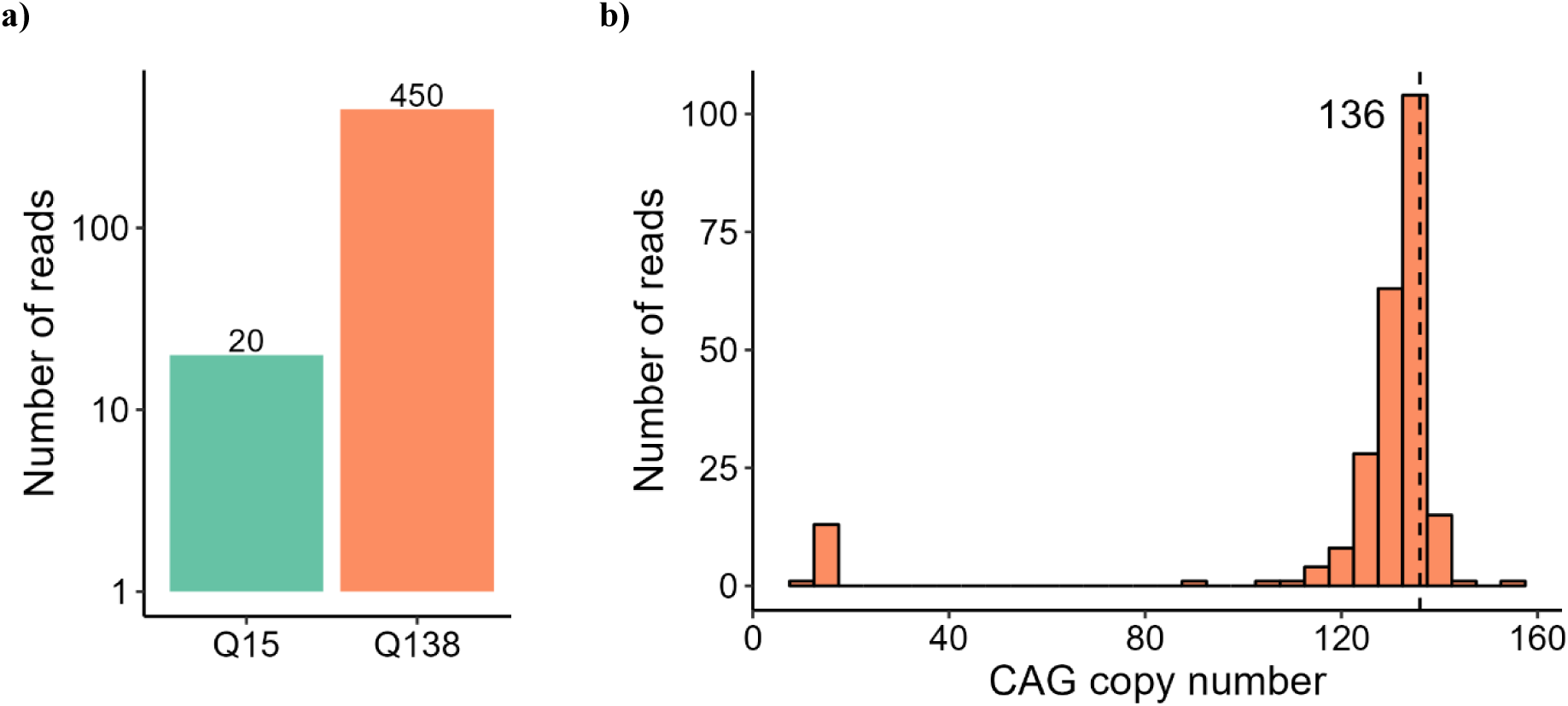
The analysis of the reads obtained by Nanopore sequencing of the pSB*tet*-Neo-HttQ15/138. Only reads containing the polyQ tract are shown. a) The number of reads corresponding to the N-terminal region of the Htt gene in the pSB*tet*-Neo-HttQ15/138 plasmids. b) The calculated CAG copy number in the reads that contained the Htt N-terminal region. The modal value of 136 CAG repeats in pSB*tet*-Neo-HttQ138 plasmid is indicated by vertical dashed line.

**Fig. S4.**
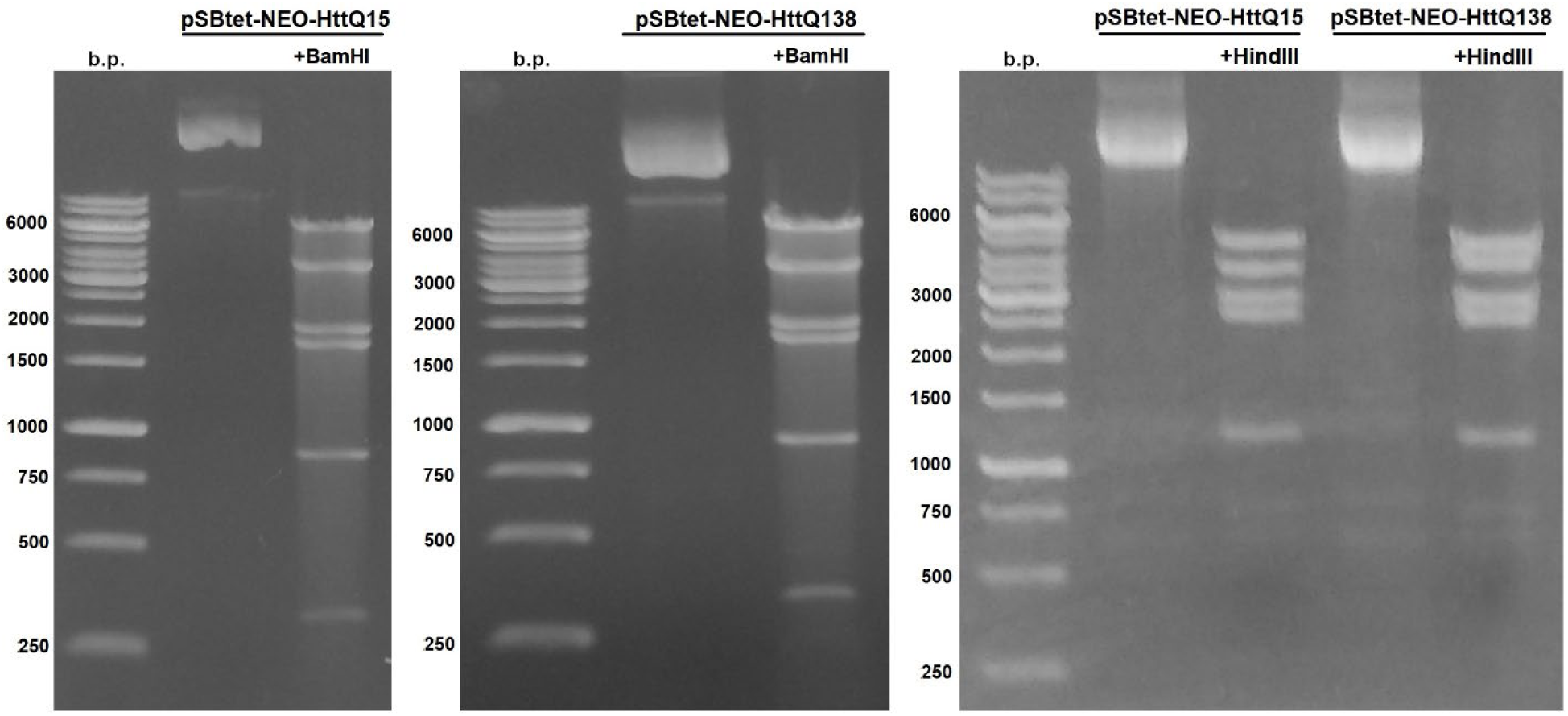
Left: agarose electrophoresis of DNA fragments after hydrolysis of pSB*tet*-Neo-Q15 or pSB*tet*-Neo-Q138 plasmids by BamHI restriction endonuclease. Right: agarose electrophoresis of DNA fragments after hydrolysis of pSB*tet*-Neo-Q15 or pSB*tet*-Neo-Q138 plasmids by HindIII restriction endonuclease.

**Fig. S5.**
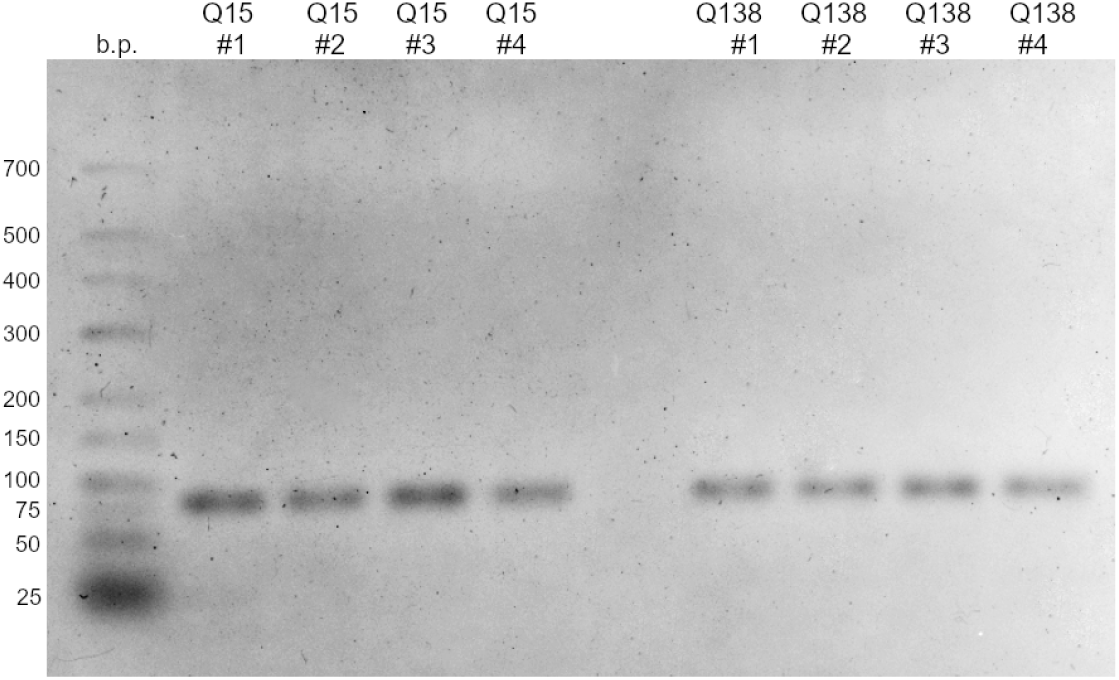
Agarose electrophoresis of amplicons obtained after qPCR analysis of monoclonal transgenic Neuro-2a lines.

**Fig. S6.**
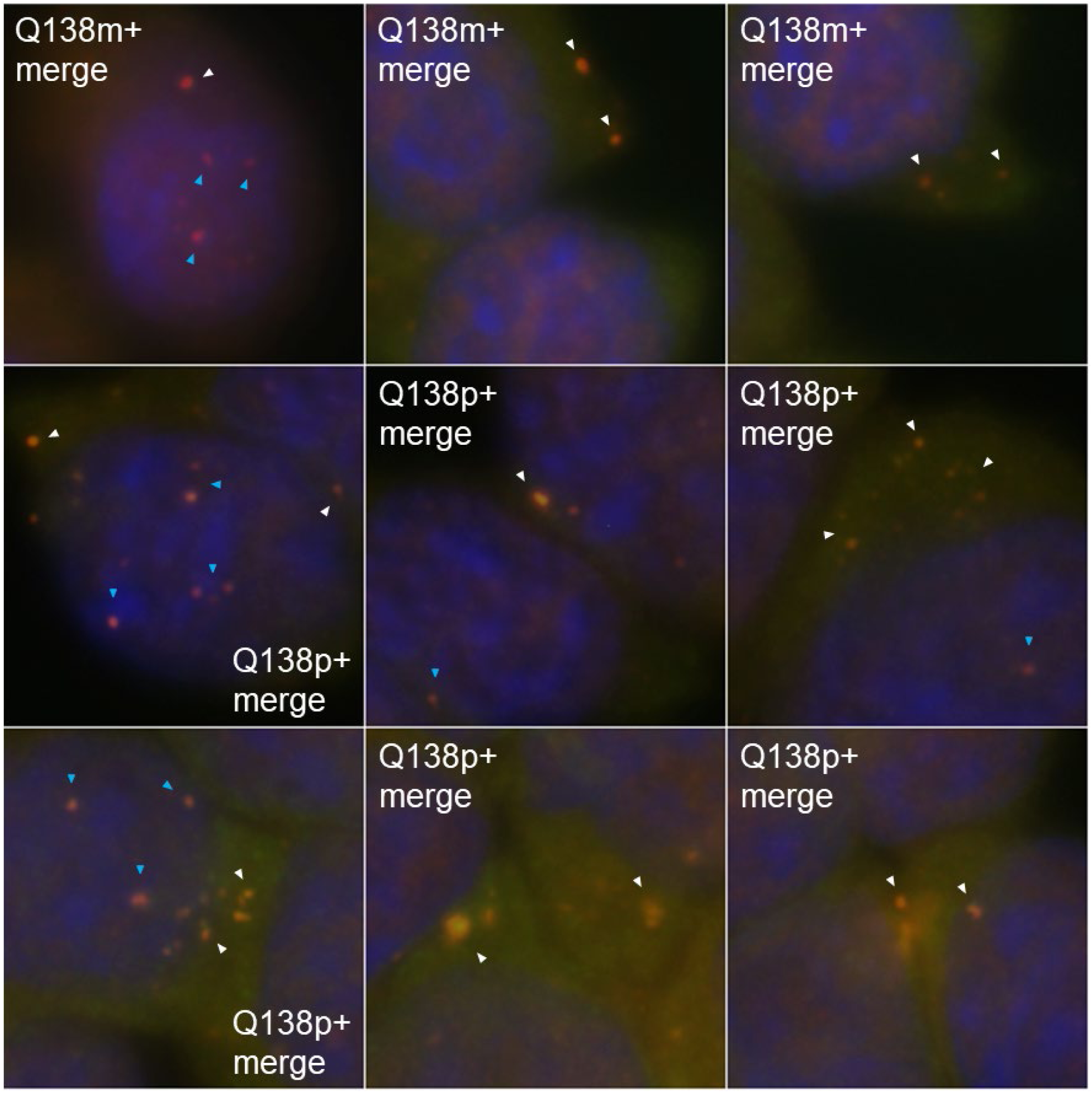
Close view of HttQ138 aggregates after 14 days of Htt expression in monoclonal and polyclonal HttQ138-expressing Neuro-2a cell lines. Htt-immunopositive inclusion bodies in Q138m+/Q138p+ cells are indicated by arrows; intranuclear inclusion bodies are indicated in blue arrows.

**Table S1.**
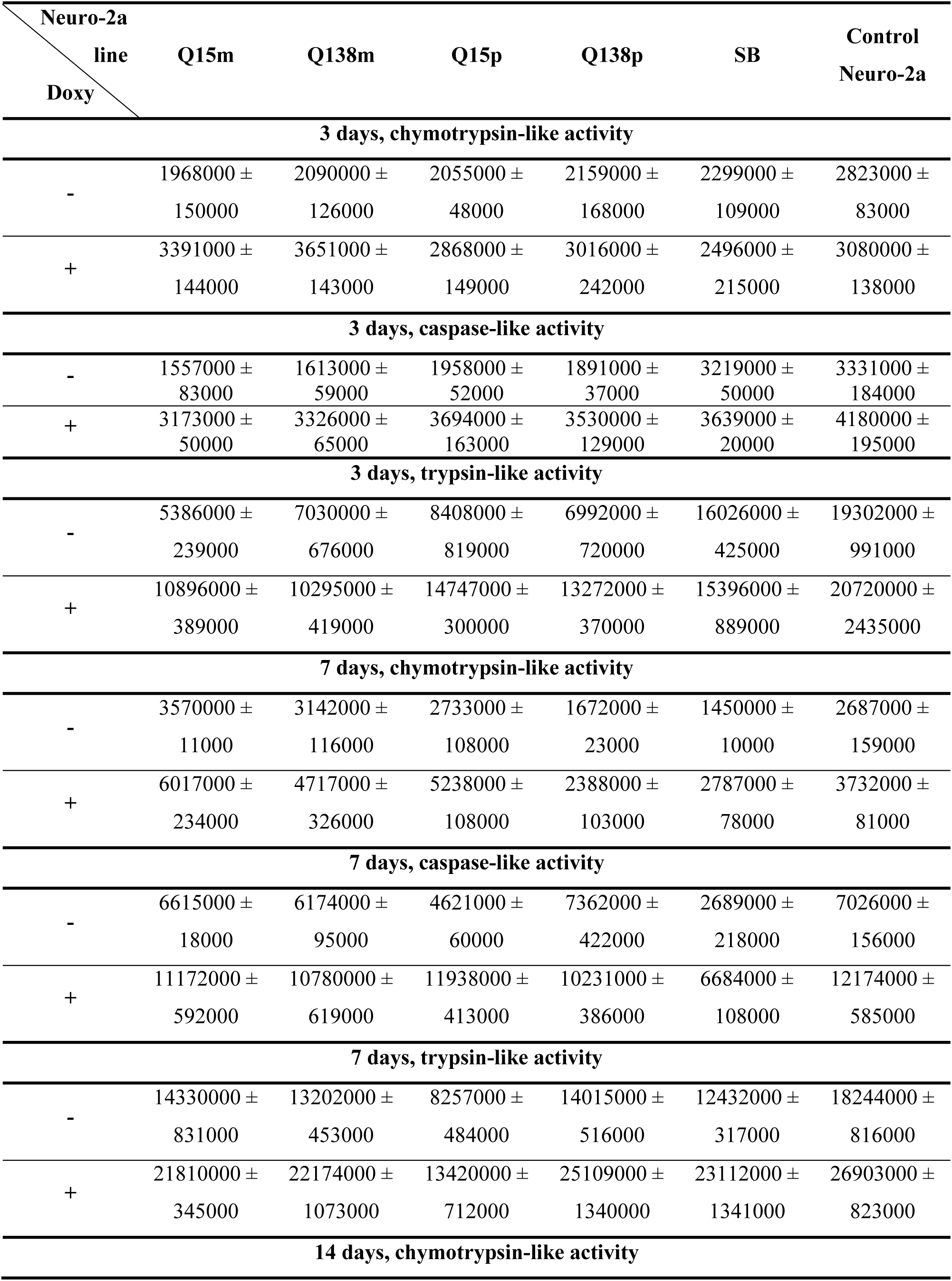
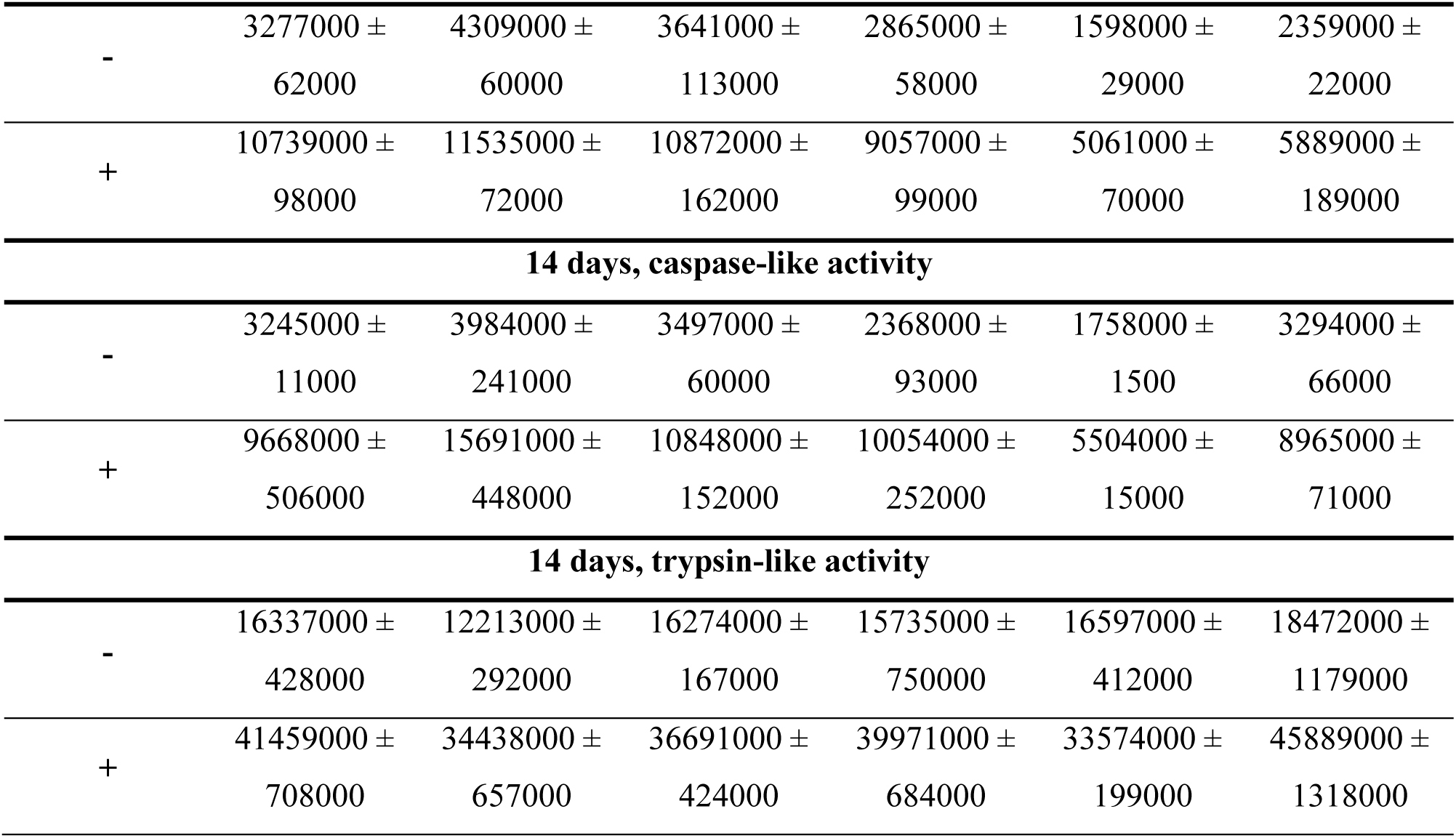
Absolute values of proteasome activity different types, U (background protease activity was subtracted) without/with expression induction of huntingtin variants in transgenic Neuro-2a lines, as well as in control Htt-non-expressing lines.

**Fig. S7.**
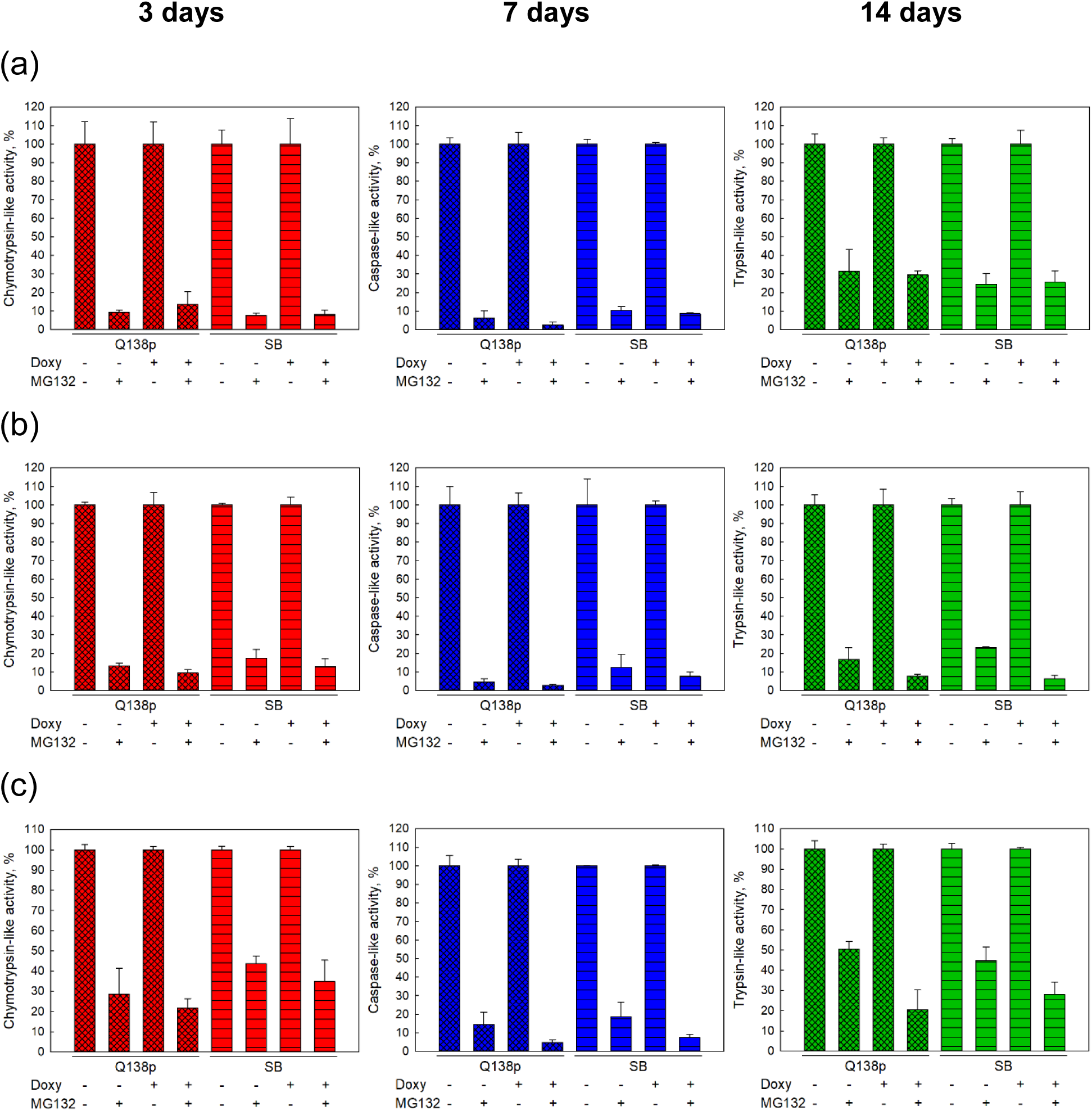
Background protease activity in transgenic Neuro-2a polyclonal HttQ138-expressing cell line and in control line (SB) without/with doxycycline induction, as well as in the absence/presence of proteasome inhibitor MG132. Chymotrypsin-like *(a)*, caspase-like *(b)* and trypsin-like *(c)* activities were evaluated. Activity assay was carried out 3 (left panel), 7 (middle panel) and 14 (right panel) days after induction. Protease activity in the MG132-non-threated cell lines was referred to 100% in each experiment.

**Fig. S8.**
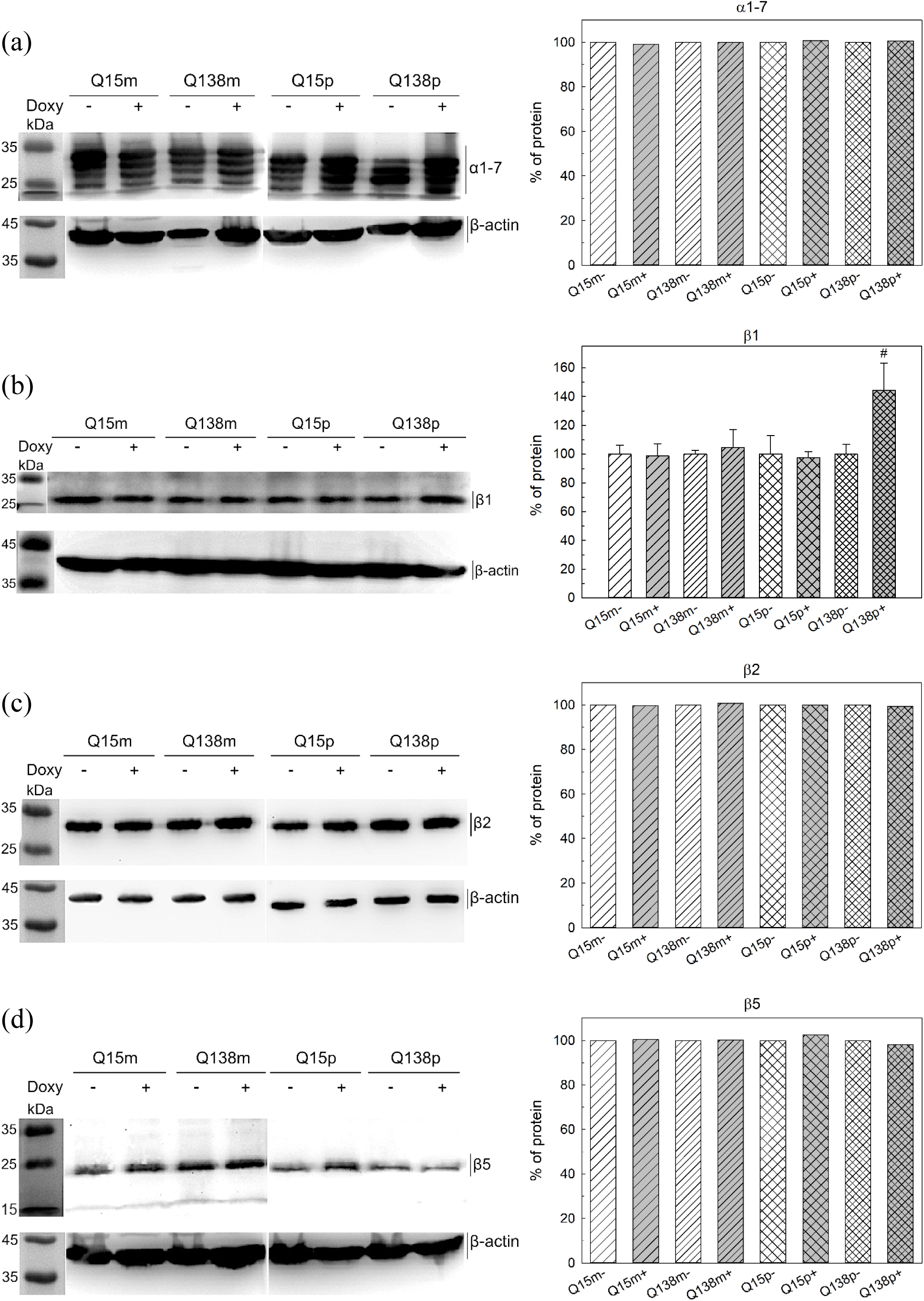
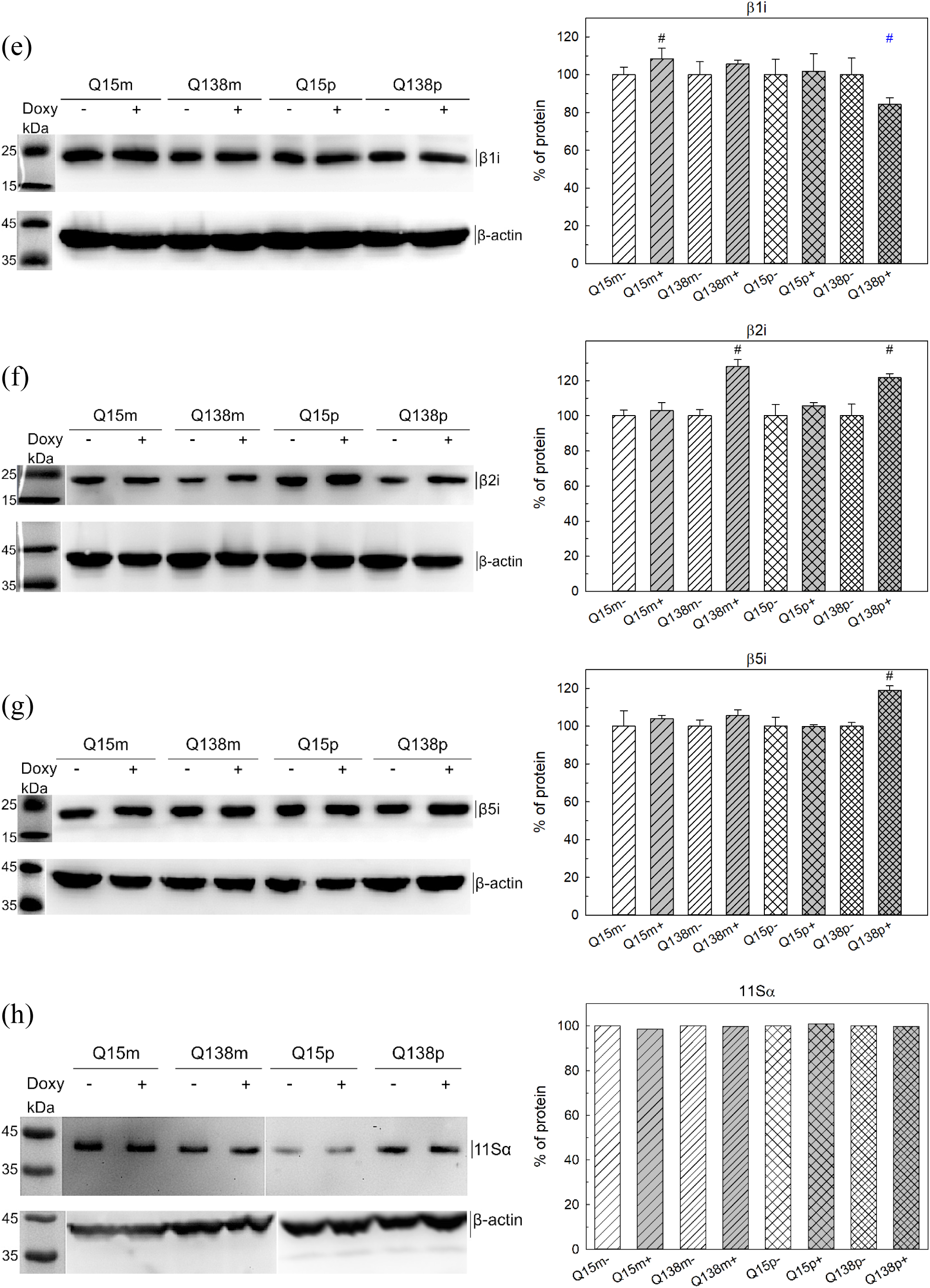
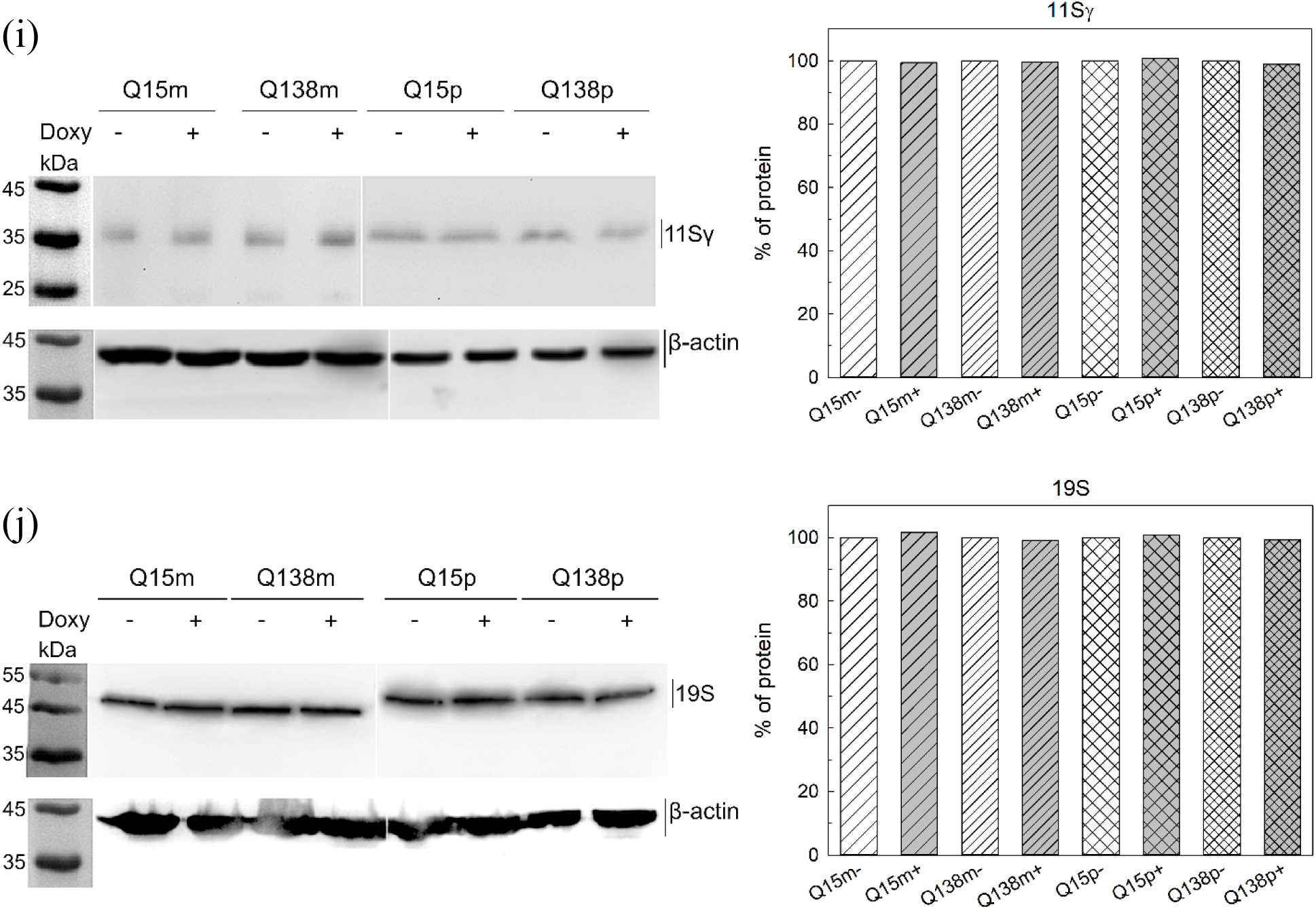
Left panel: western blotting of Neuro-2a transgenic cells lysates after 3 days of HttQ15/HttQ138 overexpression (+) and without expression (-). p, m – poly- and monoclonal cultures, respectively. All proteasomal α-subunits (α1, 2, 3, 5, 6, 7) (a), β1, β2 and β5 proteasomal subunits (b-d), β1i, β2i and β5i proteasomal immune subunits (e-g), 11Sα (h), 11Sγ (i), and 19S (j) proteins are visualized. Right panel: relative amounts of tested proteasomal subunits, 11Sα, 11Sγ and 19S proteins in transgenic Neuro-2a cells lysates after normalization on β-actin content. Protein content in the doxycycline non-threated cells was referred to 100% in each experiment. ^#^*p* < 0,05, −Doxy vs +Doxy. Blue symbols indicate *p* < 0,1.

