## Supplementary Table S1 for "Neuronal cell line expressing full-length mutant huntingtin displays alteration of proteasome activity"

**Table S1.** Absolute values of proteasome activity different types, U (background protease activity was subtracted) without/with expression induction of huntingtin variants in transgenic Neuro-2a lines, as well as in control Htt-non-expressing lines.

| **Neuro-2a line**  **Doxy** | **Q15m** | **Q138m** | **Q15p** | **Q138p** | **SB** | **Control Neuro-2a** |
| --- | --- | --- | --- | --- | --- | --- |
| **3 days, chymotrypsin-like activity** | | | | | | |
| **-** | 1968000 ± 150000 | 2090000 ± 126000 | 2055000 ± 48000 | 2159000 ± 168000 | 2299000 ± 109000 | 2823000 ± 83000 |
| **+** | 3391000 ± 144000 | 3651000 ± 143000 | 2868000 ± 149000 | 3016000 ± 242000 | 2496000 ± 215000 | 3080000 ± 138000 |
| **3 days, caspase-like activity** | | | | | | |
| **-** | 1557000 ± 83000 | 1613000 ± 59000 | 1958000 ± 52000 | 1891000 ± 37000 | 3219000 ± 50000 | 3331000 ± 184000 |
| **+** | 3173000 ± 50000 | 3326000 ± 65000 | 3694000 ± 163000 | 3530000 ± 129000 | 3639000 ± 20000 | 4180000 ± 195000 |
| **3 days, trypsin-like activity** | | | | | | |
| **-** | 5386000 ± 239000 | 7030000 ± 676000 | 8408000 ± 819000 | 6992000 ± 720000 | 16026000 ± 425000 | 19302000 ± 991000 |
| **+** | 10896000 ± 389000 | 10295000 ± 419000 | 14747000 ± 300000 | 13272000 ± 370000 | 15396000 ± 889000 | 20720000 ± 2435000 |
| **7 days, chymotrypsin-like activity** | | | | | | |
| **-** | 3570000 ± 11000 | 3142000 ± 116000 | 2733000 ± 108000 | 1672000 ± 23000 | 1450000 ± 10000 | 2687000 ± 159000 |
| **+** | 6017000 ± 234000 | 4717000 ± 326000 | 5238000 ± 108000 | 2388000 ± 103000 | 2787000 ± 78000 | 3732000 ± 81000 |
| **7 days, caspase-like activity** | | | | | | |
| **-** | 6615000 ± 18000 | 6174000 ± 95000 | 4621000 ± 60000 | 7362000 ± 422000 | 2689000 ± 218000 | 7026000 ± 156000 |
| **+** | 11172000 ± 592000 | 10780000 ± 619000 | 11938000 ± 413000 | 10231000 ± 386000 | 6684000 ± 108000 | 12174000 ± 585000 |
| **7 days, trypsin-like activity** | | | | | | |
| **-** | 14330000 ± 831000 | 13202000 ± 453000 | 8257000 ± 484000 | 14015000 ± 516000 | 12432000 ± 317000 | 18244000 ± 816000 |
| **+** | 21810000 ± 345000 | 22174000 ± 1073000 | 13420000 ± 712000 | 25109000 ± 1340000 | 23112000 ± 1341000 | 26903000 ± 823000 |
| **14 days, chymotrypsin-like activity** | | | | | | |
| **-** | 3277000 ± 62000 | 4309000 ± 60000 | 3641000 ± 113000 | 2865000 ± 58000 | 1598000 ± 29000 | 2359000 ± 22000 |
| **+** | 10739000 ± 98000 | 11535000 ± 72000 | 10872000 ± 162000 | 9057000 ± 99000 | 5061000 ± 70000 | 5889000 ± 189000 |
| **14 days, caspase-like activity** | | | | | | |
| **-** | 3245000 ± 11000 | 3984000 ± 241000 | 3497000 ± 60000 | 2368000 ± 93000 | 1758000 ± 1500 | 3294000 ± 66000 |
| **+** | 9668000 ± 506000 | 15691000 ± 448000 | 10848000 ± 152000 | 10054000 ± 252000 | 5504000 ± 15000 | 8965000 ± 71000 |
| **14 days, trypsin-like activity** | | | | | | |
| **-** | 16337000 ± 428000 | 12213000 ± 292000 | 16274000 ± 167000 | 15735000 ± 750000 | 16597000 ± 412000 | 18472000 ± 1179000 |
| **+** | 41459000 ± 708000 | 34438000 ± 657000 | 36691000 ± 424000 | 39971000 ± 684000 | 33574000 ± 199000 | 45889000 ± 1318000 |
