## Supplementary Figures S1-S8 for "Neuronal cell line expressing full-length mutant huntingtin displays alteration of proteasome activity"

**Supplementary Materials**

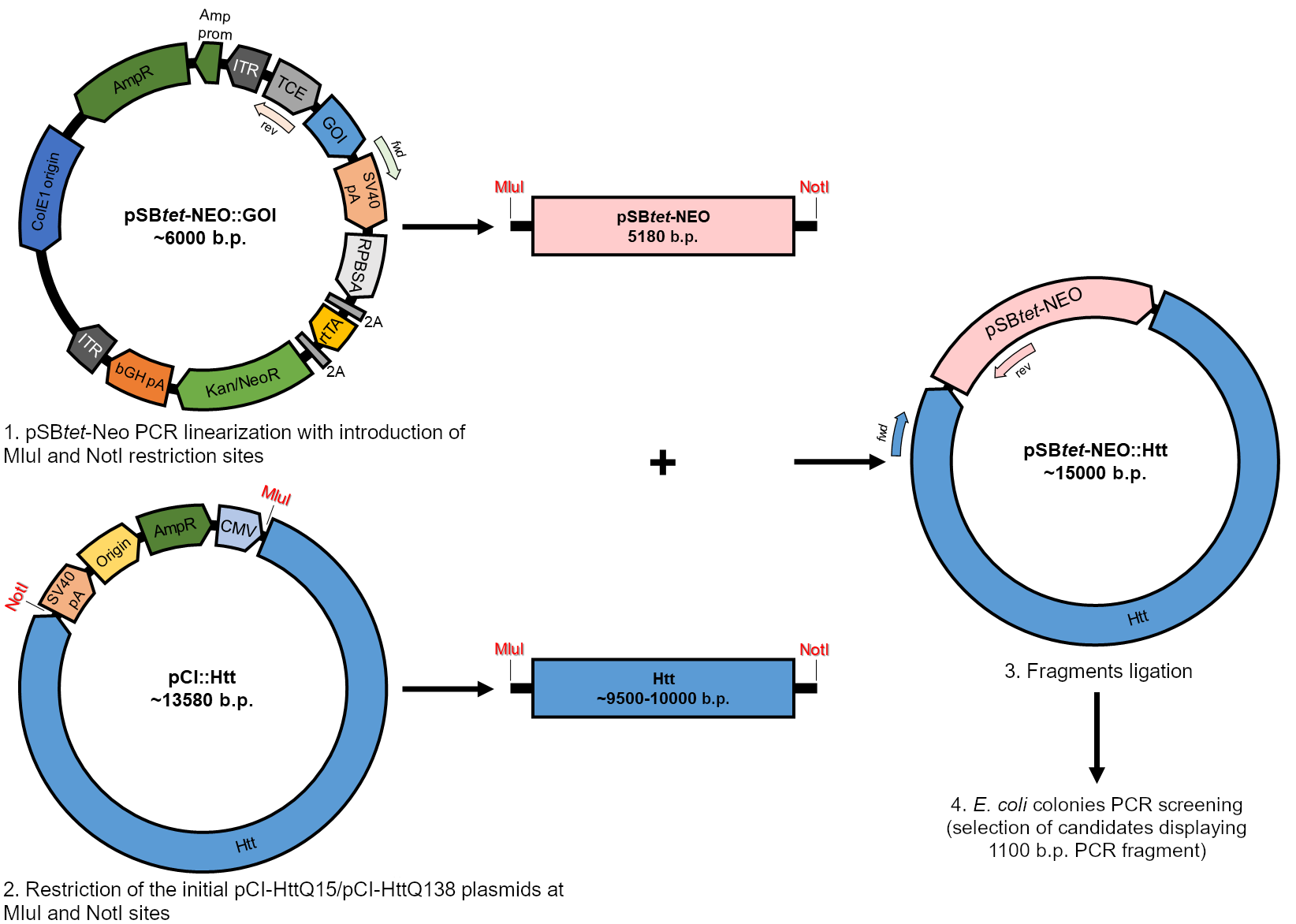

**Fig. S1.** Scheme of Htt genes cloning into the pSB*tet*-Neo expression vector. ITR — inverse terminal repeats, TCE — tetracycline-inducible promoter, GOI — studied gene, RPBSA — synthetic constitutive promoter, 2A — flanking fragments with proteolysis sites, rtTA — tetracycline-inducible transactivator, Kan/NeoR, AmpR — kanamycin/neomycin and ampicillin resistance genes, respectively, SV40 pA and bGH pA — polyadenylation signals, CMV — CMV immediately-early enhancer/promoter, Htt — normal or mutant huntingtin gene.

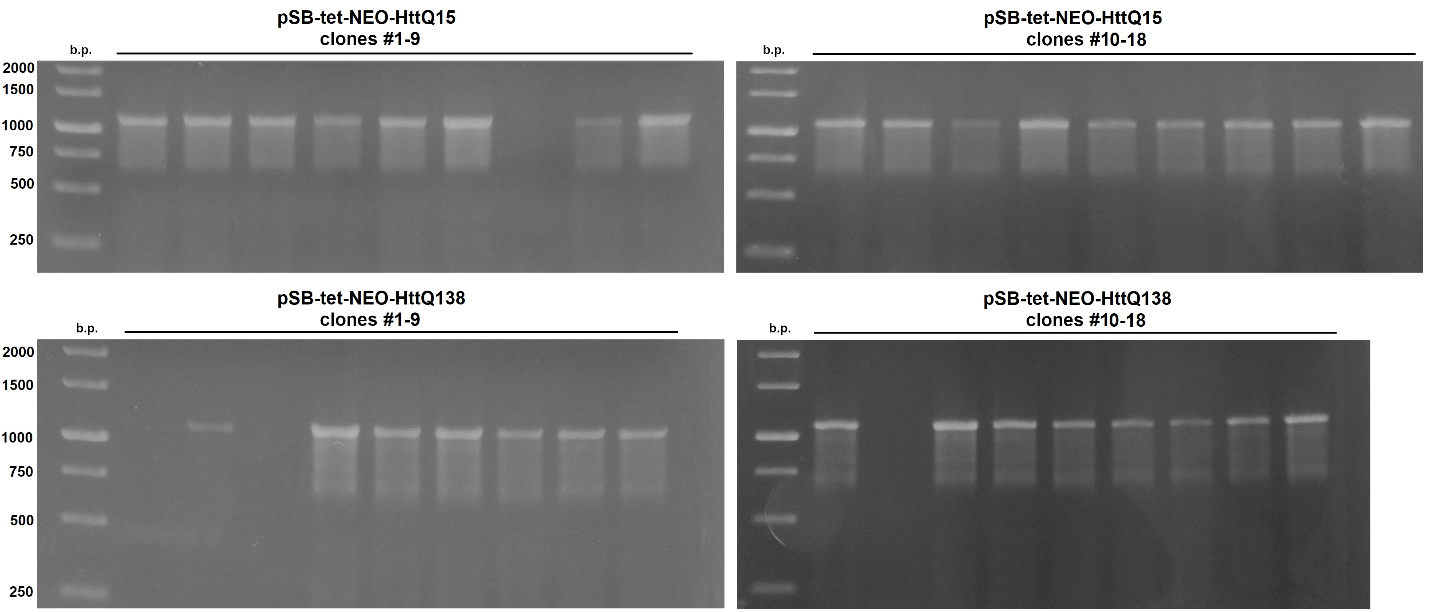

**Fig. S2.** Agarose electrophoresis of PCR products obtained during the PCR selection of *E. coli* X1-Blue clones containing the pSB*tet*-Neo-HttQ15 or pSB*tet*-Neo-HttQ138 constructs.

| **a)**  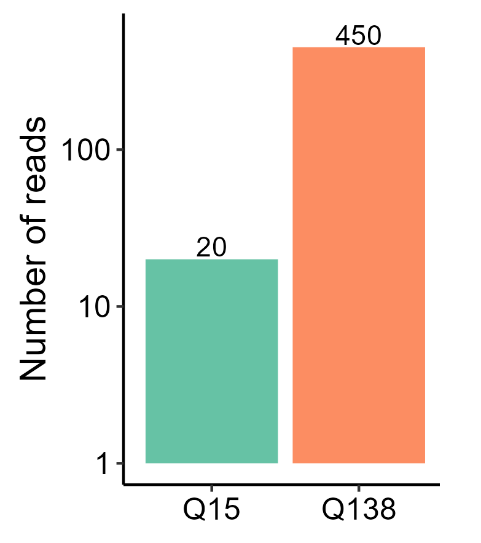 | **b)**  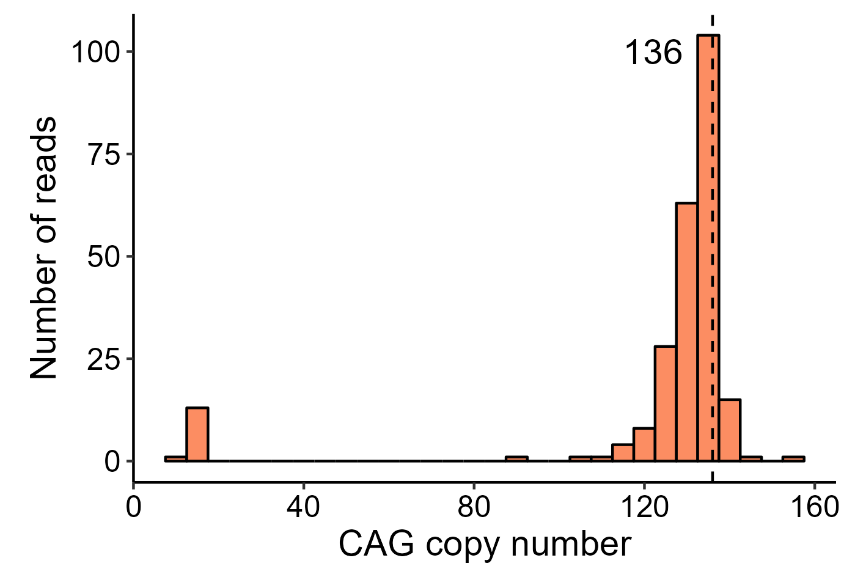 |
| --- | --- |

**Figure S3.** The analysis of the reads obtained by Nanopore sequencing of the pSB*tet*-Neo-HttQ15/138. Only reads containing the polyQ tract are shown. a) The number of reads corresponding to the N-terminal region of the Htt gene in the pSB*tet*-Neo-HttQ15/138 plasmids. b) The calculated CAG copy number in the reads that contained the Htt N-terminal region. The modal value of 136 CAG repeats in pSB*tet*-Neo-HttQ138 plasmid is indicated by vertical dashed line.

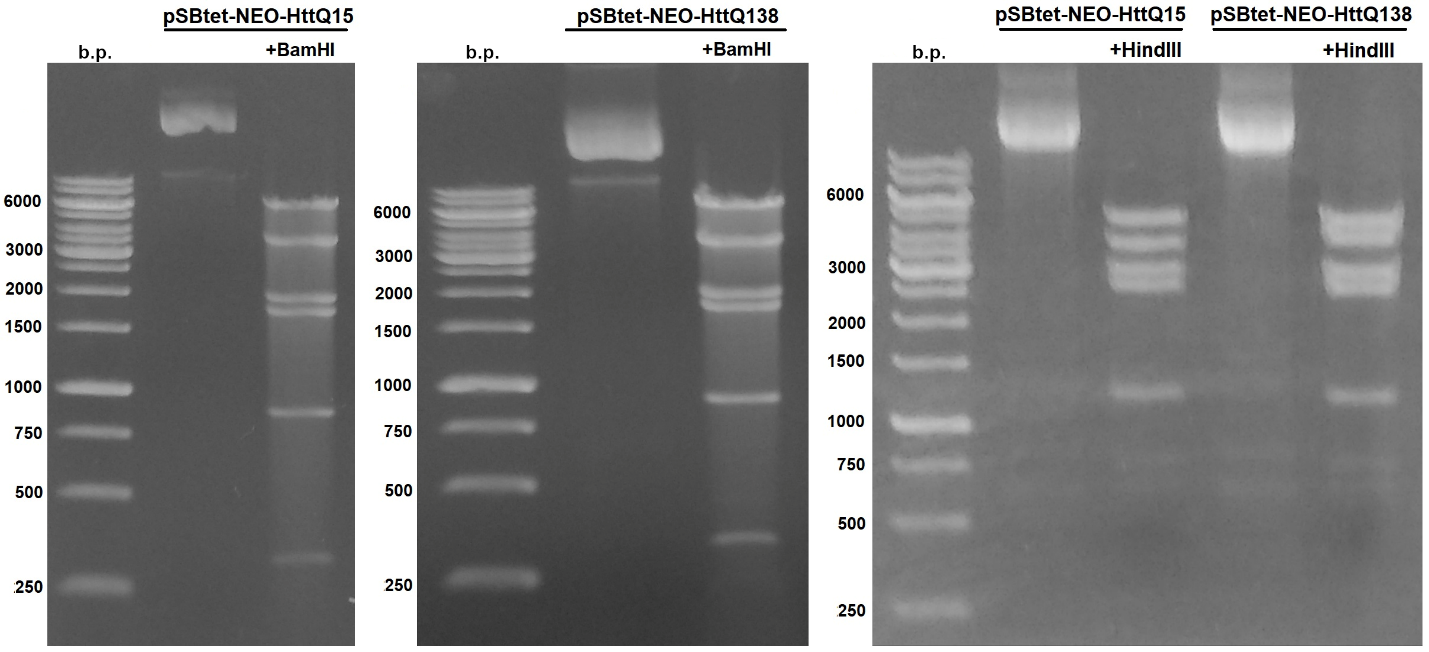

**Fig. S4.** Left: agarose electrophoresis of DNA fragments after hydrolysis of pSB*tet*-Neo-Q15 or pSB*tet*-Neo-Q138 plasmids by BamHI restriction endonuclease. Right: agarose electrophoresis of DNA fragments after hydrolysis of pSB*tet*-Neo-Q15 or pSB*tet*-Neo-Q138 plasmids by HindIII restriction endonuclease.

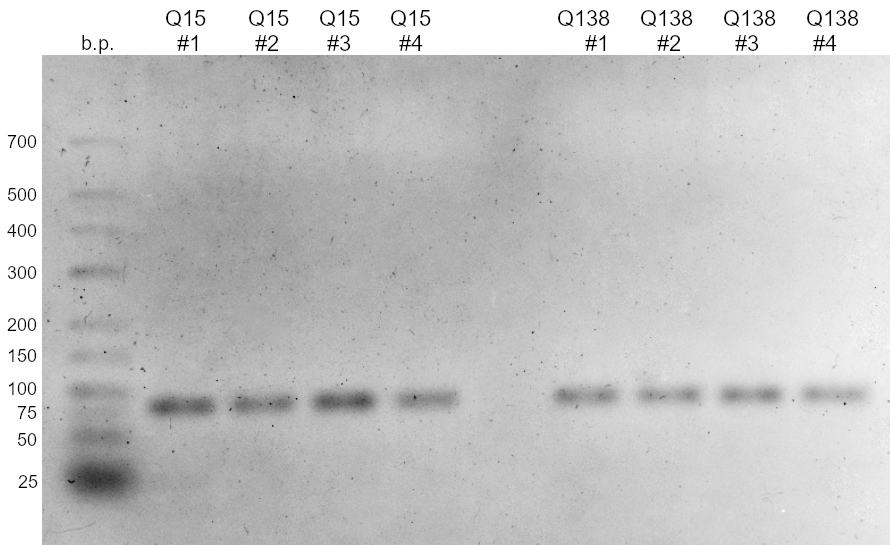

**Fig. S5.** Agarose electrophoresis of amplicons obtained after qPCR analysis of monoclonal transgenic Neuro-2a lines.

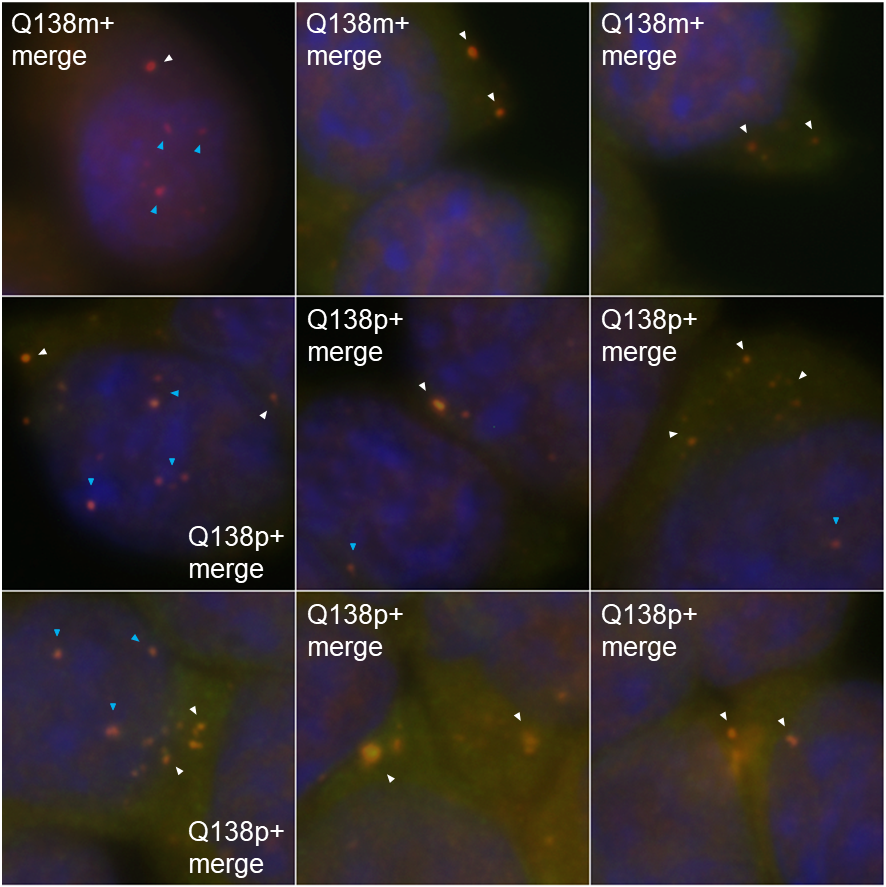

**Fig. S6.** Close view of HttQ138 aggregates after 14 days of Htt expression in monoclonal and polyclonal HttQ138-expressing Neuro-2a cell lines. Htt-immunopositive inclusion bodies in Q138m+/Q138p+ cells are indicated by arrows; intranuclear inclusion bodies are indicated in blue arrows.

| **3 days** | **7 days** | **14 days** |
| --- | --- | --- |
| (a)  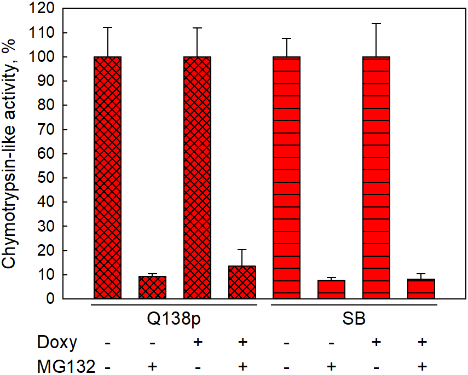 | 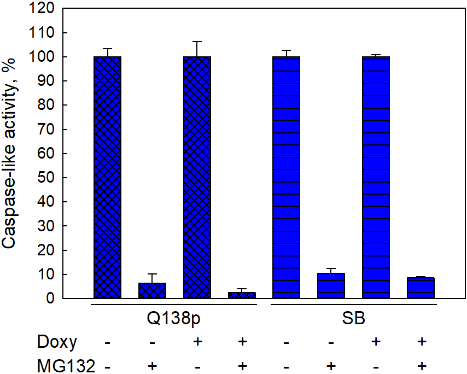 | 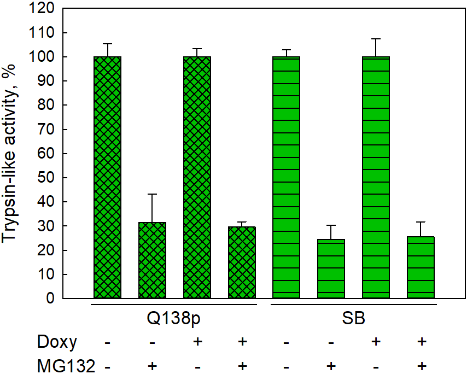 |
| (b)  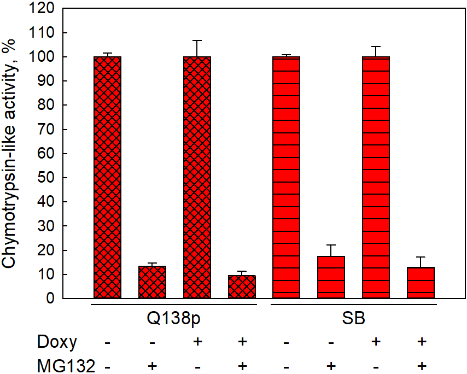 | 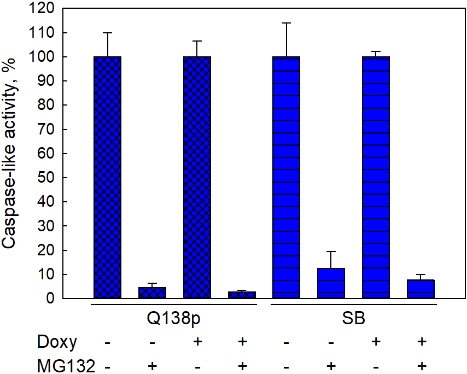 | 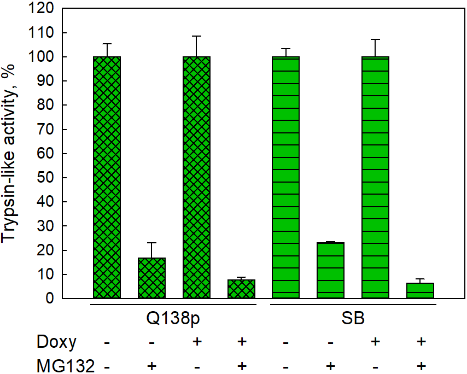 |
| (c)  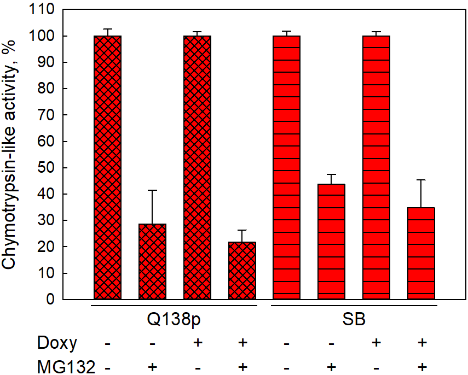 | 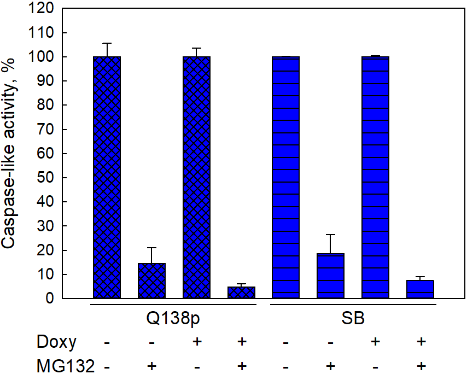 | 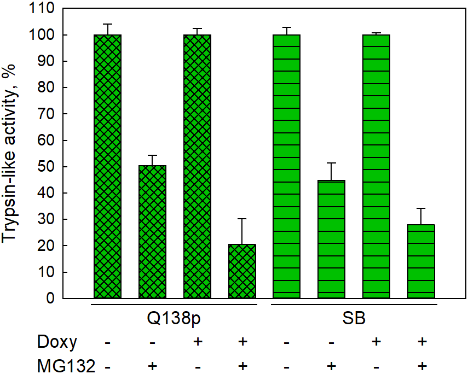 |

**Fig. S7.** Background protease activity in transgenic Neuro-2a polyclonal HttQ138-expressing cell line and in control line (SB) without/with doxycycline induction, as well as in the absence/presence of proteasome inhibitor MG132. Chymotrypsin-like *(a)*, caspase-like *(b)* and trypsin-like *(c)* activities were evaluated. Activity assay was carried out 3 (left panel), 7 (middle panel) and 14 (right panel) days after induction. Protease activity in the MG132-non-threated cell lines was referred to 100% in each experiment.

| (a)  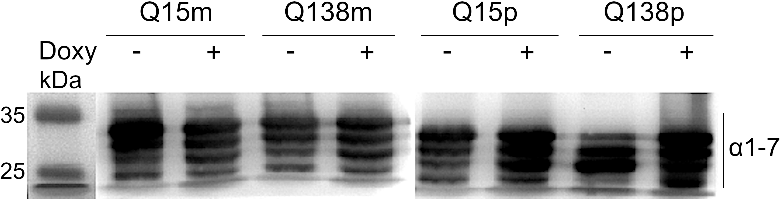  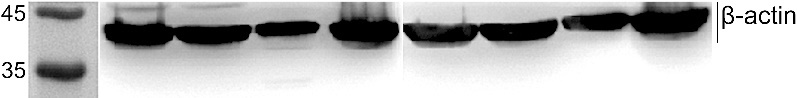 | 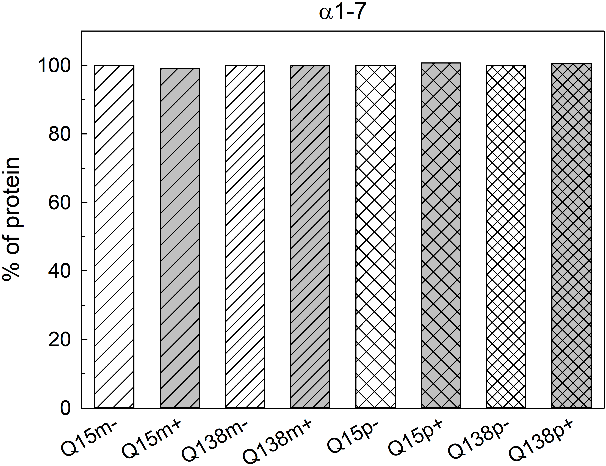 |
| --- | --- |
| (b)  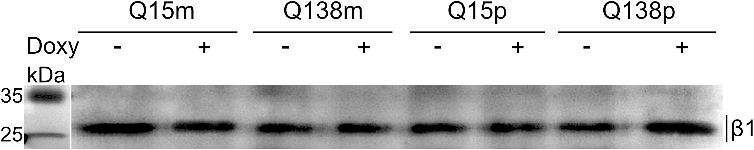  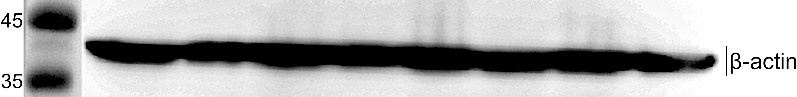 | 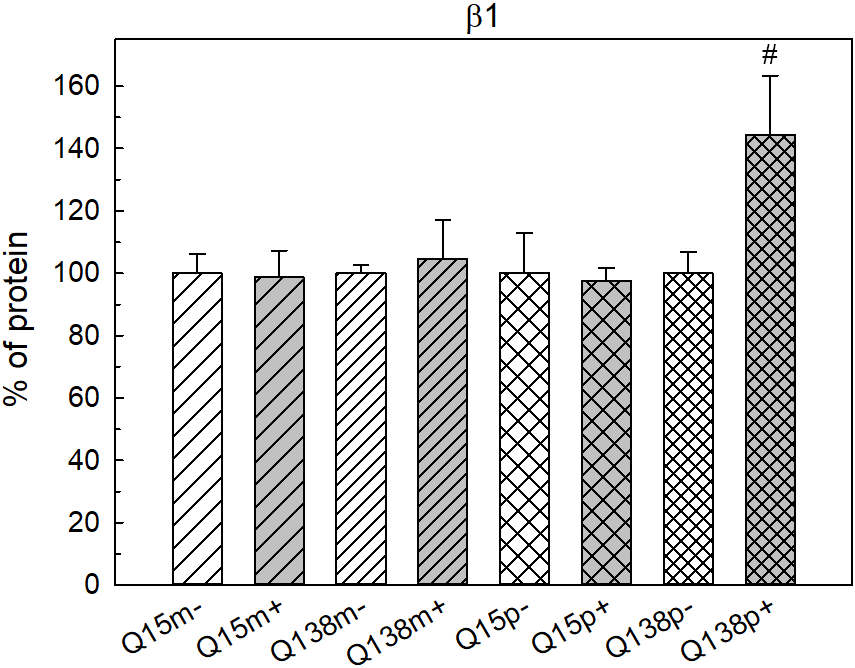 |
| (c)  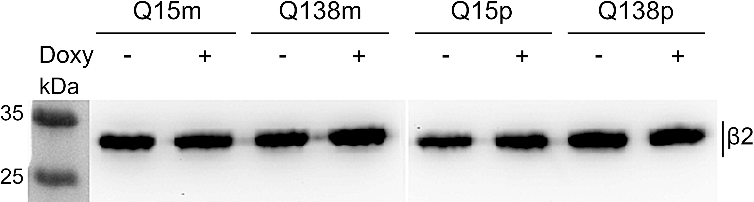  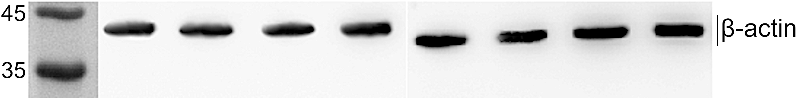 | 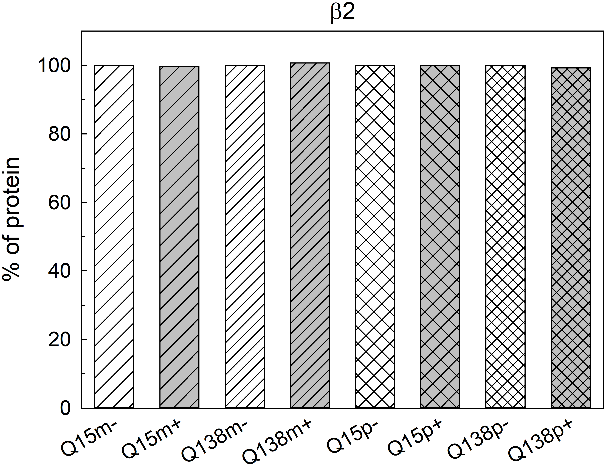 |
| (d)  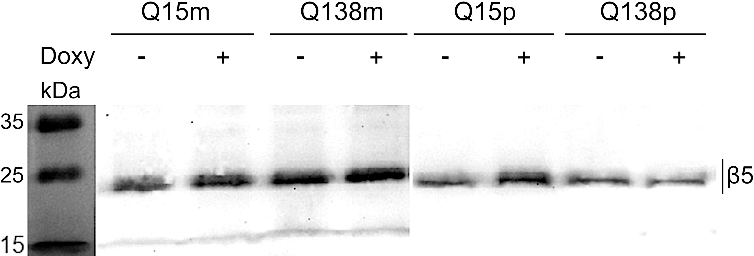  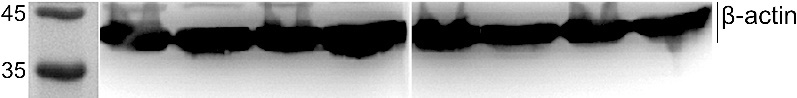 | 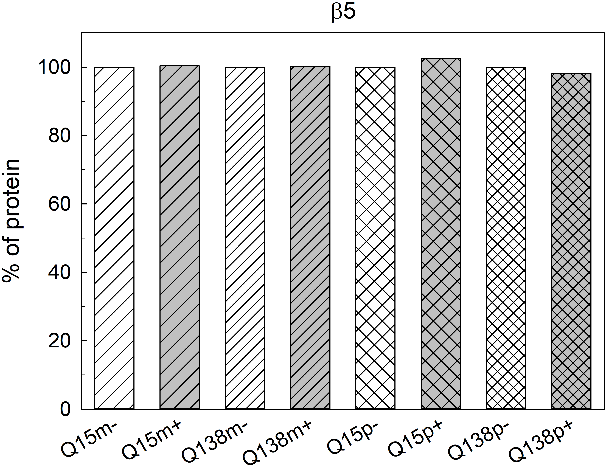 |
| (e)  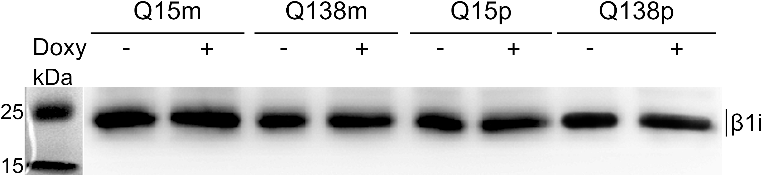  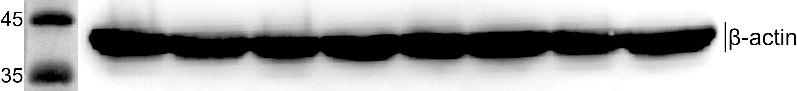 | 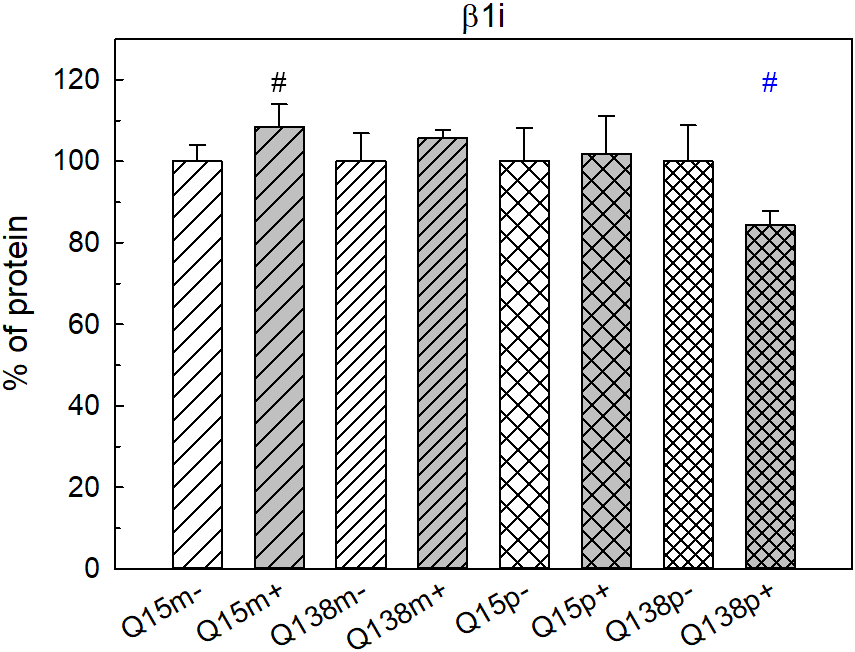 |
| (f)  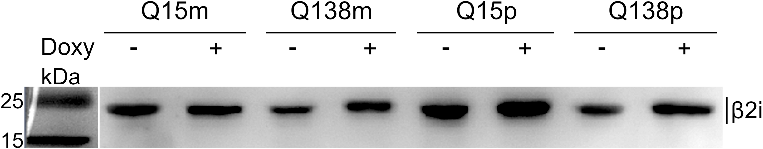  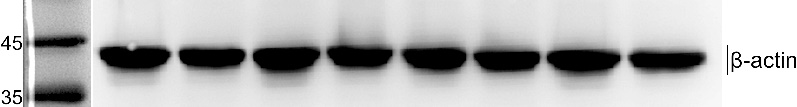 | 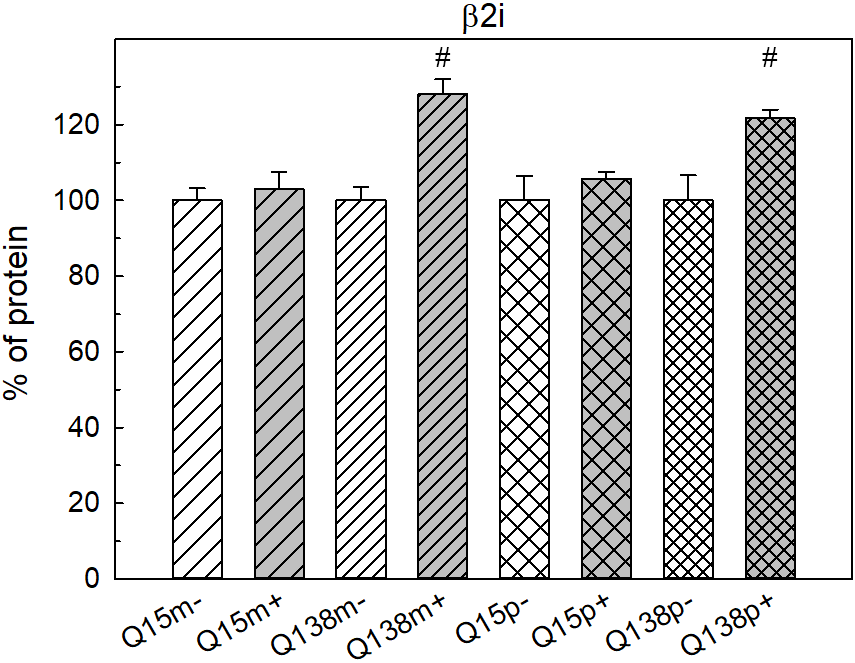 |
| (g)  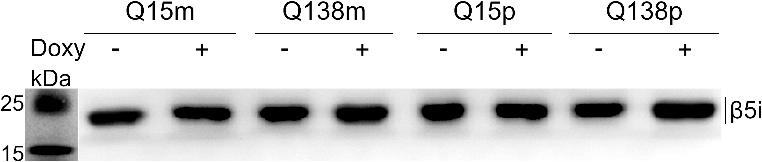  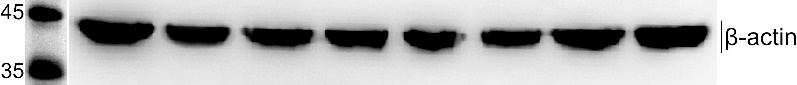 | 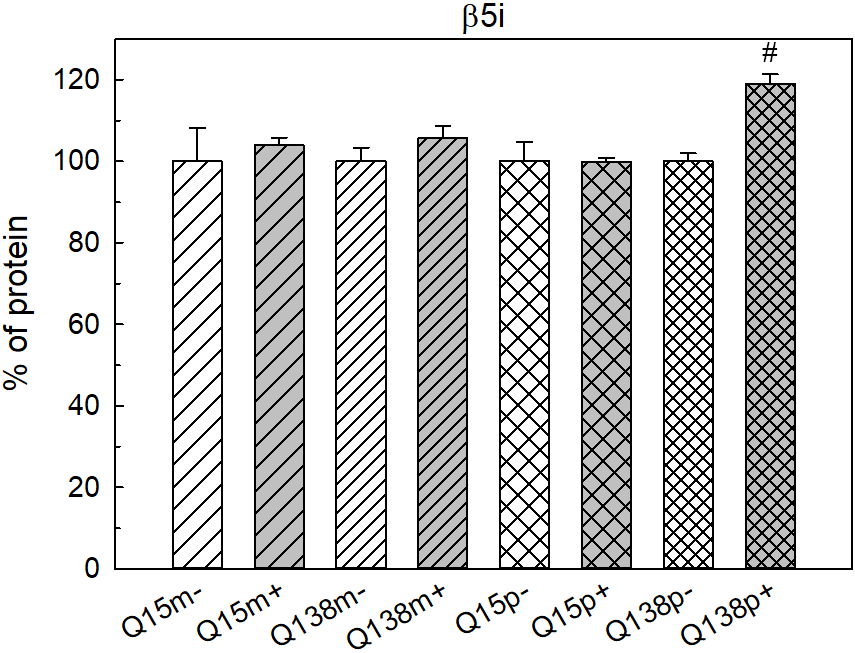 |
| (h)  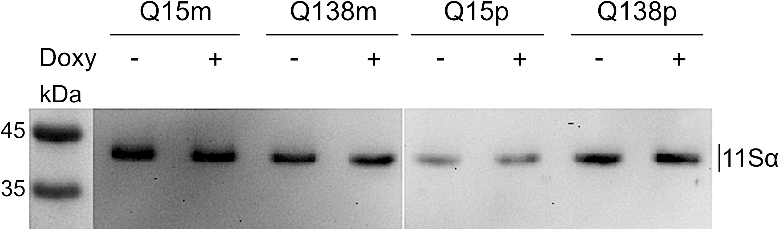  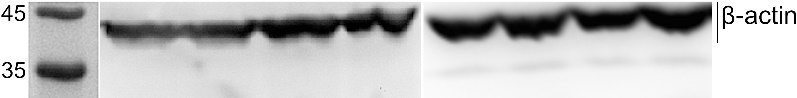 | 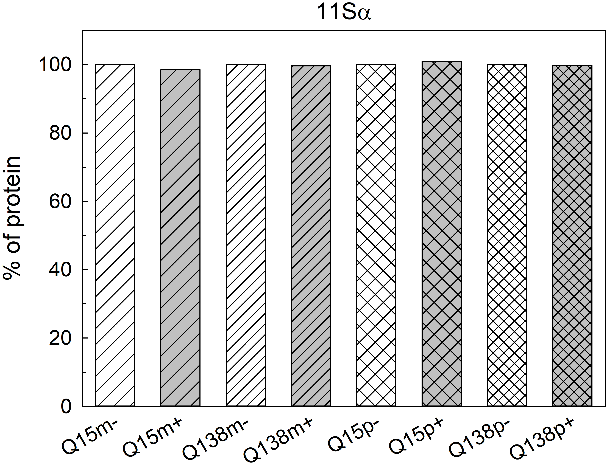 |
| (i)  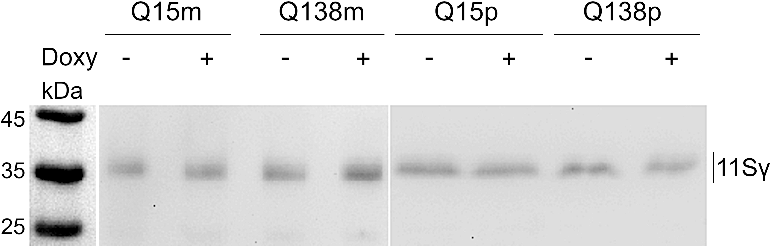  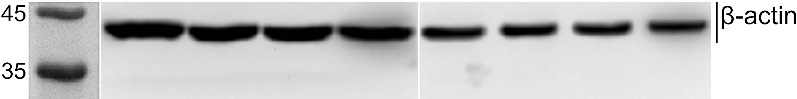 | 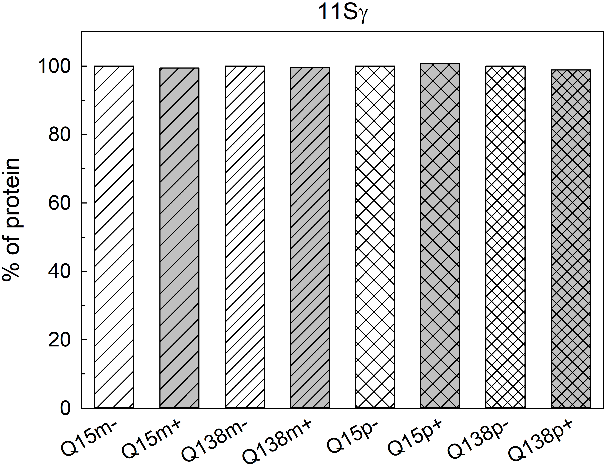 |
| (j)  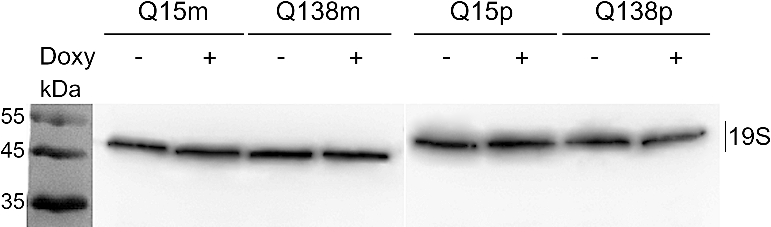  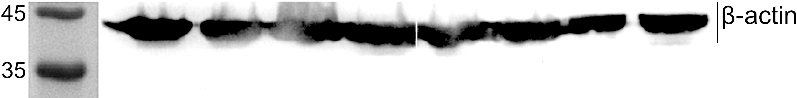 | 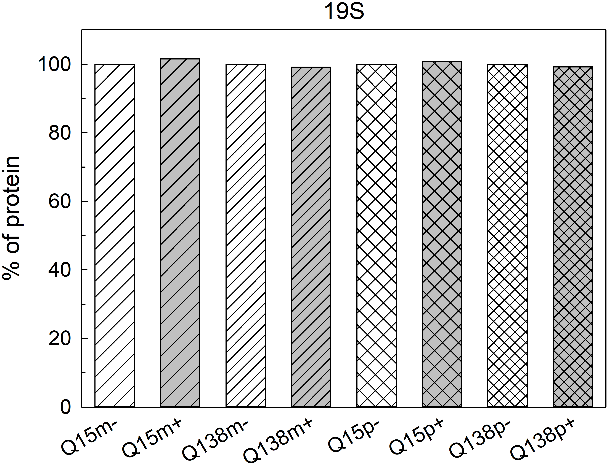 |

**Fig. S8.** Left panel: western blotting of Neuro-2a transgenic cells lysates after 3 days of HttQ15/HttQ138 overexpression (+) and without expression (-). p, m – poly- and monoclonal cultures, respectively. All proteasomal α-subunits (α1, 2, 3, 5, 6, 7) (a), β1, β2 and β5 proteasomal subunits (b-d), β1i, β2i and β5i proteasomal immune subunits (e-g), 11Sα (h), 11Sγ (i), and 19S (j) proteins are visualized. Right panel: relative amounts of tested proteasomal subunits, 11Sα, 11Sγ and 19S proteins in transgenic Neuro-2a cells lysates after normalization on β-actin content. Protein content in the doxycycline non-threated cells was referred to 100% in each experiment. ^#^*p* < 0,05, -Doxy vs +Doxy. Blue symbols indicate *p* < 0,1.
